# The high quality Chinese white truffle genome and novel fossil-calibrated estimate of Pezizomycetes divergence reveal the tempo and mode of true truffles genome evolution

**DOI:** 10.1101/2024.11.26.625401

**Authors:** Jacopo Martelossi, Jacopo Vujovic, Yue Huang, Alessia Tatti, Kaiwei Xu, Federico Puliga, Yuanxue Chen, Omar Rota Stabelli, Fabrizio Ghiselli, Xiaoping Zhang, Alessandra Zambonelli

## Abstract

The genus *Tuber* (family: Tuberaceae) includes the most economically valuable ectomycorrhizal (ECM), truffle-forming fungi. Previous genomic analyses revealed that massive transposable element (TE) proliferation represents a convergent genomic feature of mycorrhizal fungi, including Tuberaceae. Repetitive sequences are one of the major drivers of genome evolution shaping its architecture and regulatory networks. In this context, Tuberaceae represent an important model system to study their genomic impact; however, the family lacks high-quality assemblies. Here, we tested the interplay between TEs and Tuberaceae genome evolution by producing a highly contiguous assembly for the endangered Chinese truffle *Tuber panzhihuanense*, along with a novel timeline for Tuberaceae diversification and comprehensive comparative genomic analyses. We found that concurrently with a Paleogene diversification of the family, pre-existing Chromoviridae-related Gypsy clades independently expand in different truffle lineages leading to increased genome size and high gene family turnover rates, but without resulting in highly scrambled genomes. Additionally, we found an enrichment of ECM-induced gene families among ancestral duplication events. Finally, we explored the repetitive structure of nuclear ribosomal DNA (rDNA) loci for the first time in the clade. We found that most of the 45S rDNA paralogues are undergoing concerted evolution, though an isolated divergent locus raises concerns about potential issues for metabarcoding and biodiversity assessments. Our study provides a fundamental genomic resource for future research on truffle genomics and showcases a clear example on how establishment and self-perpetuating expansion of heterochromatin can drive massive genome size variation due to activity of selfish genetic elements.

## Introduction

Fungi play a pivotal role in the functioning of ecosystems (Boddy et al. 2016) representing core components of nutrients cycling, carbon cycling, and likely carbon sequestration (Mehar and Sundaramoorthy 2018; Emilia Hannula and Morriёn 2022). Nowadays nearly 90% of land plants are found to be in association with one or more fungal partners and fungi have probably played a crucial role in the plant colonisation of land around ∼500 million years ago (Bonfante 2003). The hyphae of mycorrhizal fungi establish a strict relationship with plant roots facilitating the acquisition by the plant of water and nutrients such as phosphorus and nitrogen (Martin et al. 2016).

Based on the type of association established with their host, mycorrhizal fungi are usually subdivided in different groups: ectomycorrhizal (ECM), arbuscular mycorrhiza, orchid mycorrhiza, and ericoid mycorrhiza (Brundrett and Tedersoo 2018). Truffles are characterised by a hypogeous fruit body where spores are sequestered, they typically establish ECM relationships with plants (Tedersoo et al. 2010) and rely on pungent aromas to attract the animals responsible for their dispersal (Bonito et al. 2013). Similarly to other symbiotic fungi, truffle-forming species have independently diversified dozens of times during fungi evolution, both within Ascomycota and Basidiomycota (Tedersoo et al. 2010). Many truffle species are edible for humans and because of their aroma they have been considered as food delicacies for centuries (Mello et al. 2006). The Tuberaceae family (Acomycota: Pezizomycetes) is the most rich and diverse clade of truffle-forming fungi and among them the *Tuber* genus (true truffles) contains the most economically valuable species (Leonardi et al. 2021), such as the Périgord black truffle *Tuber melanosporum* Vittad. and the italian white truffle *Tuber magnatum* Pico. Most of the host plants of Tuberaceae are angiosperms and presumably these plants were the original hosts of their most recent common ancestor with multiple independent transitions to Pinaceae in individual lineages during their evolution (Bonito et al. 2013).

*T. melanosporum* is the first *Tuber* to have its genome sequenced (Martin et al. 2010), revealing one of the most complex fungal genome sequenced to date, characterised by a 4-fold larger genome compared to other ascomycetes, a high transposable element (TE) content, low gene redundancy, and a restricted set of genes encoding for carbohydrate cleaving enzymes. Comparative genomic analyses across Pezizomycetes (Murat et al. 2018) and dozens of other mycorrhizal-forming fungi (Miyauchi et al. 2020) revealed that these are common genomic features of Tuberaceae as well as other symbiotic fungi highlighting also the importance of secreted proteins (SPs), lineage-restricted genes, and gene co-option in the establishment of an ECM lifestyle. To control TE activity *T. melanosporum*, relies on a non-exhaustive, partly reversible methylation process, known as the methylation-induced premeiotically (MIP) defence system (Montanini et al. 2014). MIP is more similar to TE control systems of metazoans and plants rather than of those of other fungi, such as *Neurospora crassa*, which rely on an highly efficient repeat-induced point mutation system (RIP; Galagan and Selker 2004). Indeed, *T. melanosporum* MIP is highly selective with hypermethylation of TE rich genomic regions and hypomethylation of TEs in close proximity to genes.

Transposable elements and other repetitive sequences are known to be one of the major causes of structural variations within individuals and species, promoting gene duplication, gene loss, genomic rearrangements and reshaping the overall genomic regulatory network (Bourque et al. 2018). In this context, the highly repetitive Tuberaceae genome represents an important system to study the genomic impact of transposons and their possible role in affecting phenotypic traits. However, due to their repetitiveness, TEs also impose challenges in genome assembly especially when relying only on short read data, leading to highly fragmented genomes and unreliable gene and transposon annotations (Peona et al. 2021; Rhie et al. 2021).

Here, we studied the highly repetitive nature of Tuberaceae genomes by sequencing the Chinese white truffle *Tuber panzhihuanense* X.J. Deng & Y. Wang using the PacBio HiFi technology. Along with *Tuber latisporum* Juan Chen & P.G, from which it was recently splitted based on barcoding analyses (Deng et al. 2013), *T. panzhihuanense* is considered one of the most economically important truffle species in China (Wan et al. 2015). Despite its potential economic value, it is currently listed as critically endangered species by the Redlist of China’s Biodiversity — Macrofungi (www.mee.gov.cn; Yao et al. 2020; Zhuang et al. 2020) and is widely agreed that it cannot be cultivated yet. Our highly contiguous assembly represents a substantial improvement compared to previously available Pezizales genomes allowing us to characterise and study in unprecedented detail the impact of TE bursts in Tuberaceae genome evolution. Additionally, we produced a novel genome-based timeline for the diversification of the family—using the largest number of fossils to date for the divergence time estimation of an Ascomycota—which placed the dawn of the *Tuber* genus during the Paleogene. Using these novel genomic resources we show that increased Tuberaceae genome size is driven by lineage-specific, Gypsy-mediated expansion of heterochromatic genomic regions, resulting in more than half of the genome composed by ∼96% of TEs. These massive TE bursts did not lead to scrambled genome architectures, but impacted the evolution of multi-copy gene families. Interestingly, we also found that expanded gene families in the stem branch of Tuberaceae are more likely to be related to ECM compared to the null expectation, implying that gene duplication might have contributed to the transition from a saprotrophic to an ECM lifestyle. Finally, we describe for the first time in the clade the structure of nuclear rDNA loci, which is of particular interest due to the use of the internal transcribed spacer (ITS) as the primary barcode marker in fungi (Schoch et al. 2012). The high-quality *T. panzhihuanense* assembly represents a fundamental resource for future agriculture-applied studies (e.g., transitioning this currently uncultivable species from wild harvesting to orchard cultivation, Lemmond et al. 2023) and will continue to serve as a basis for genomic studies of truffles.

## Results

### A high quality assembly for the Chinese white truffle

Using kmer-based approaches, we estimated a genome size of 110 Mb, compatible with estimates of other *Tuber* species (Murat et al. 2018). As expected for the haploid state of the gleba fruiting body tissue, the genome resulted completely homozygous (Supplementary Figure S1A). The assembled genome spans 119 Mb with a high N50 value of 7.3 Mb, a consensus quality value (QV) of 57.72 and complete single copy BUSCO scores of 96.4%. (Ascomycota odb_10; Supplementary Table S1; Supplementary Figure S1B). The assembly consists of 29 contigs with contigs 1 to 17 representing nearly the whole nuclear genome (Fig. 1A) on which we predicted 8,701 protein coding genes (PCGs) with a genome-wide PCG density of 0.12. On the predicted proteome we obtained a complete and single copy BUSCO score slightly higher than the assembly score. (Supplementary Table S1). The *T. panzhihuanense* genome encodes for 363 secreted proteins (SP) of which 173 are small SP (SSPs) and a reduced set of 198 carbohydrate-active enzymes (CAZys; Supplementary Table S2). The contig 18 represents a 302 kb mitochondrial genome for which we obtained a complete and circularised representation.

**Fig 1:**
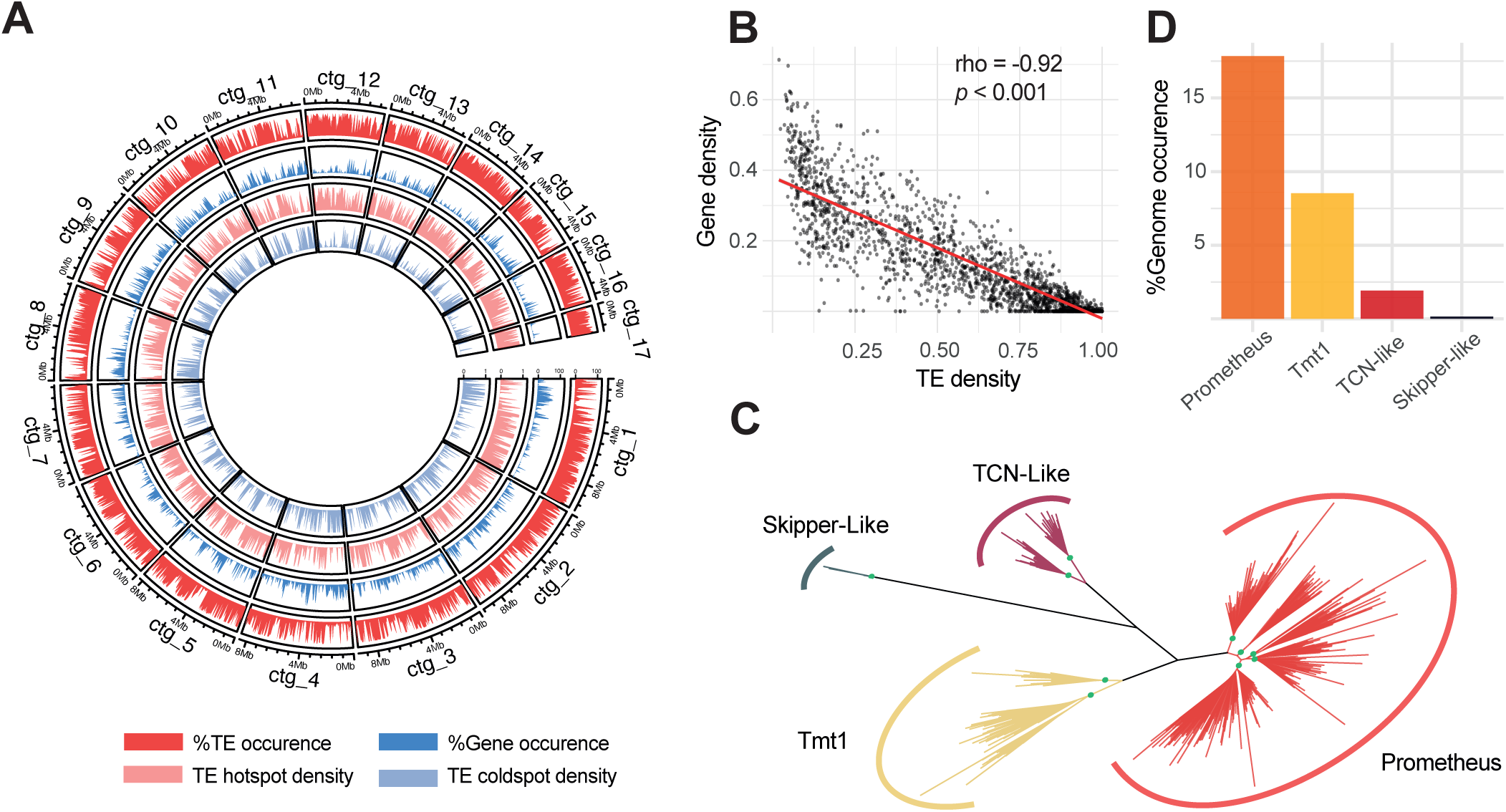
A highly compartmentalised genome colonised by Chromoviridae-related Gypsy transposons. **(A)** From outer to inner circles: distribution of transposons, genes, TE hotspots, and TE coldspots across the 17 main contigs of the *T. panzhihuanense* nuclear genome. **(B)** Correlation between TE density against gene density across 50 kb non-overlapping genomic windows; rho = Spearman’s rank correlation coefficient. **(C)** Phylogenetic tree of representative Gypsy RT protein segments longer than 100 amino acids. Green circles highlight crown nodes of identified families with bootstrap support values ≥ 75. **(D)** Genomic occurrence of the four identified Gypsy clades in the *T. panzhihuanense* genome.

Compared to previously published Pezizales genomes available on NCBI, our assembly has a N50 value 2.4 times greater compared to the scaffold N50 of the second most contiguous genome (*Morchella crassipes* (Vent.) Pers.; GCA_009192285.1) and 4 times greater when considering only true truffles (*Tuber magnatum* Picco; GCA_003182015.1).

### A highly compartmentalised genome dominated by retrotransposons

To improve TE annotation in a reasonable time frame (Goubert et al. 2022; Peona et al. 2024), we manually curated the automatically generated libraries, selecting the most represented consensus sequences and/or those with detectable protein fragments. Although we curated only 18% of the raw library, we found that curated TEs accounted for 54% of the total TE content, demonstrating that our selection strategy successfully captured most of the TE complement. While the total TE content and its composition did not change much when using the raw versus the partially curated TE library, resulting in both instances over 60%, we observed a drastic reduction in the number of unclassified elements using the former (supplementary figure S2 A-B). Retrotransposons, and particularly Gypsy long terminal repeats (LTRs) and long interspersed nuclear element (LINE) Tad1 elements, greatly dominate the genomic TE landscape of the Chinese white truffle, covering 26.6% and 17% of the genome, while DNA and rolling circle (RC) transposons cover 9.9% and 0.6%, respectively.

The genomic distribution of TEs and genes highlights a highly compartmentalised genomic organisation with a strong and significant negative correlation between TE and gene density (Spearman’s rho = -0.92; *P* < 0.01; Fig. 1A-B). The majority of the assembly is composed of either significantly TE-enriched (TE hotspots; 51% of the genome; mean TE content = 96%) or TE-depleted genomic regions (TE coldspots; 32% of the genome; mean TE content = 15%; supplementary table S3) with TE hotspots significantly longer than TE coldspots (Welch Two Sample t-test, *P* < 0.01; supplementary figure S3). The relative proportions of the 3 main transposon groups (Gypsy, LINEs, and DNA transposons) to the total amount of genomic space are different between the two genomic compartments with long retrotransposons being predominant on TE hotspots and DNA elements in TE coldspots (supplementary figure S4). Moreover, TE coldspots contain almost all PCGs (7,047), with a gene density of 0.32, 2.6 times higher than the genome wide estimation, while only 354 genes are located in TE-rich genomic regions. These results are mirrored by the distance of TEs from the closest gene (supplementary figure S5 A), with DNA transposons more closely associated with genes compared to LINEs and LTRs (Kruskal-Wallis rank sum test, *P* < 0.01; Pairwise Wilcoxon rank-sum test with Bonferroni correction, *P* < 0.01), likely due to the existence of numerous short non-autonomous and/or highly degenerated versions (supplementary figure S5 B-C).

When looking at the genomic distribution of effector-like SPs, we did not identify an enrichment among genes contained within TE hotspots (sample size = 8701; *X*^2^ = 1.6749 , *p* = 0.25). The compartmentalization of transposons in true truffle genomes appears therefore different from the frequent TE/effector compartmentalisation of pathogenic fungi genomes (Raffaele et al. 2010; Schmidt et al. 2013; Fouché et al. 2020).

### Chromoviridae are the most abundant Gypsy elements in the Chinese white truffle genome

Due to the significant contribution of Gypsy elements to the high TE content in the *T. panzhihuanense* genome, as well as in other true truffles (Payen et al. 2016; Murat et al. 2018), we further investigated their phylogenetic placement and domain structure. Network clustering identified 5 communities of Gypsy LTRs (supplementary figure S6 A), which we further split into 10 evolutionary distinct families based on phylogenetic analyses on their reverse transcriptase (RT) domain (Fig. 1 C; supplementary figure S6 B). These families belong to 4 distinct clades: Skipper-Like, Tmt1, TCN-Like, and a previously unknown but rich and diversified clade that we named Prometheus (Fig. 1C; supplementary figure S7). We were able to reconstruct a full-length consensus sequence for all but one family (Tpan_Tmt1.1; supplementary figure S8). Similarly to Tmt1 and TCN-Like elements, Prometheus LTRs possess a chromodomain (CHD) and belong to the Chromoviridae Gypsy branch (supplementary figure S8). On the other hand, the identified Skipper-like element, despite its placement with high support in a sister relationship with the reference Skipper transposon (Bootstrap = 88; Supp. Fig. supplementary figure S7) is lacking the characteristic CHD domain (Marín and Lloréns, 2000). Coherently with the high number of Prometheus transposons identified through network analyses (supplementary figure S6 A), when annotating the *T. panzhihuanense* genome with the representative sequences of each family, we found that Prometheus is the most abundant Gypsy clade (Fig. 1D; 17.71% of the genome) followed by Tmt1 (8.56%), TCN-like (1.91%), and Skipper-like (0.15%). Most of the Gypsy insertions are composed of degenerated and/or fragmented elements (supplementary figure S9), however, we also observed 3,066 insertions that almost perfectly match LTR regions of their parent consensus sequence, reassembling solo-LTR elements that arise from non-allelic homologous recombination (NAHR) between the 2 flanking LTRs (Kent et al., 2017). On the other hand, only 445 full-length copies were identified of which 29 are putative autonomous elements (supplementary table S4).

### Contrasting mode of rDNA loci evolution

Nuclear rDNA loci are some of the most challenging genomic regions to assemble because of their high repetitiveness and high homogenization of tandemly duplicated copies (Hall et al. 2022). Our high quality genome allowed us to have a first insight into rDNA structure and organisation. Indeed, self alignments of contigs 19 to 29, which are sensibly smaller compared to others (Fig. 2A) revealed the presence of a conserved pattern in their structure represented by a core sequence which includes 45S rDNA genes (18S-5.8S-28S) surrounded by a large array of tandemly repeated elements (Fig. 2B-C). We could identify other characteristic motifs: a 160 bp sequence at the 3’ end of each rDNA unit with strong similarity to LTR regions of Prometheus Gypsy transposons, whereas at the 5’ end before the main tandemly repeated region a 100 bp palindromic structure followed by a smaller GC-rich tandem repeat (Fig. 2C). Beside these 45S rDNA-only contigs, we identified other 4 complete and 2 partial 45S loci with the same structure at one end of contig 16 and contig 17 as well as an additional one isolated along contig 3, 2.8 Mb away from the closest scaffold end (Fig. 2A).

**Fig 2.**
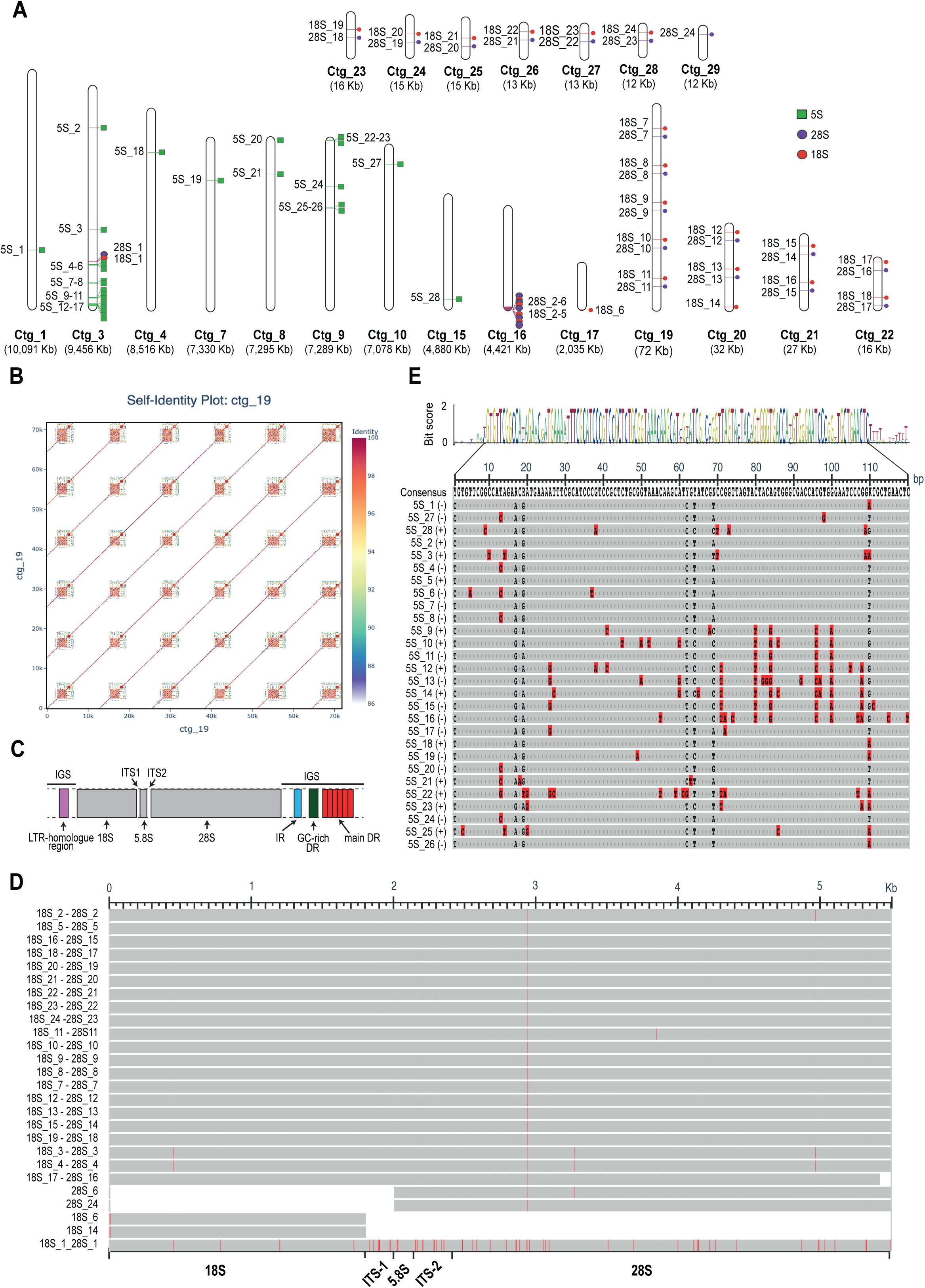
Genomic features of rDNA loci. **(A)** Genomic locations of rDNA genes. The size of contigs is proportional to their length, but different scales are used for contigs ctg_1 to ctg_17, and ctg_19 to ctg_29. The length of each contig is reported below its name. **(B)** Self-alignment of ctg_19 as an example of the 45S rDNA locus structure, as it represents the largest contig containing only 45S rDNA. **(C)** Simplified structure of 45S rDNA loci; the length of blocks is not proportional to their actual length. IR = inverted repeat, DR = direct repeat, IGS = intergenic spacer, ITS1 and ITS2 = internal transcribed spacers. **(D)** Alignment of 45S rDNA loci from 18S to 28S. Red lines highlight nucleotides that differ from the consensus sequence. The names of each locus reflect the naming system used in panel A. **(E)** Same as (D), but for 5S rDNA genes.

5S ribosomal genes were only identified outside 45S rDNA loci, with a total of 28 copies scattered along multiple contigs (Fig. 2A). We observed a tendency of 5S genes to colocalize with gene-rich and TE depleted genomic regions, with 22 copies present within TE coldspots.

We aligned all 45S rDNA complete and partial loci and all 5S rDNA genes to evaluate a possible intragenomic variation. 45S rDNA loci identified from contig 16 to contig 29 are highly homogenised, without any insertions or deletions and only 7 positions harbouring variants unique or shared across paralogue copies (Fig. 2D). On the other hand, the intergenic spacer (IGS) shows variable lengths ranging from 6,312 bp to 8,510 bp due to different copy numbers in the main tandem array (Fig. 2B). Contrary to other 45S loci, rDNA paralogue genes found isolated on contig 3 are more diverging (Fig. 2D). Compared to the 18S_10 and 28S_10 rDNA locus (arbitrarily chosen because of the high homogenization of all copies), the most diverging regions are the 2 ITS both showing 96% of identity, whereas 18S, 28S, and 5.8 show an identity of 99%, 99%, and 98% respectively. This locus is flanked by 2 IGS regions significantly shorter than those previously described (length of ∼3,600 and ∼1,600 bp; supplementary figure S10 A-B). The longer IGS is characterised by the insertion of a non-repetitive genomic region of 480 bp of unknown origin, followed by a solo-LTR (supplementary figure S9 C-D).

On the other hand, 5s rDNA genes, which are highly dispersed throughout the genome, present lower homology levels that can drop down to 80.8% of identity between 5S_14 and 5S_22 paralogues with a mean identity of 93.6% (Fig. 2E).

### A Paleogene diversification for true truffles

To study the evolution of the TE content during Tuberaceae diversification we firstly estimated a novel divergence time among a set of 32 Ascomycetes, including 8 true truffles (Fig. 3). The species were selected to maximise the number of possible calibration points, allowing us to use the highest number of fossils with clear phylogenetic placement (10 fossil calibrations) to date for an Ascomycota divergence time estimation, assembling a phylogenomic supermatrix of 1,057 complete and single-copy BUSCO genes with high occupancy (at least 95% of the species) for a total of 534,184 amino acid positions after trimming.

**Fig. 3.**
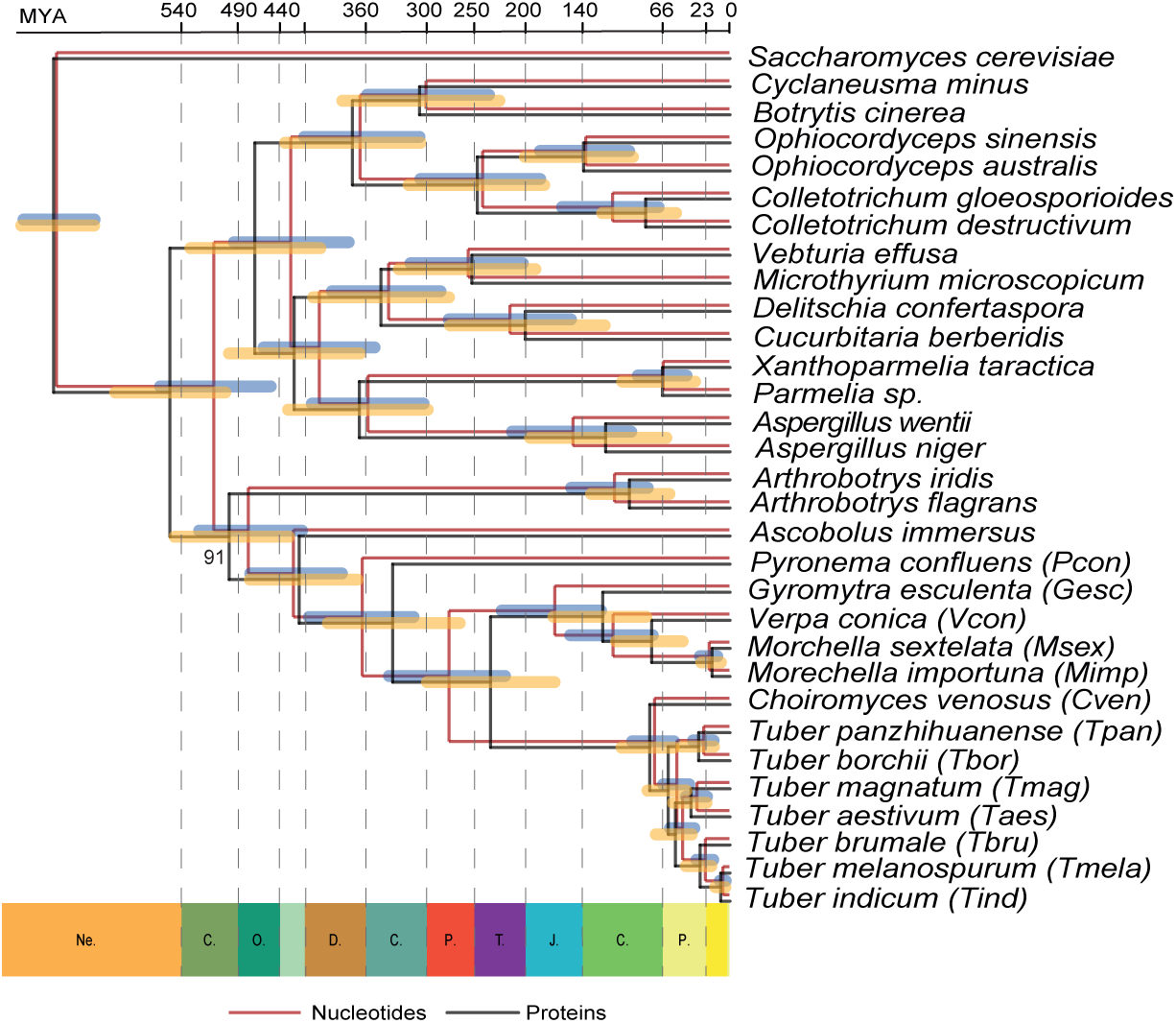
Late Cretaceous emergence of Tuberaceae and Cenozoic diversification of true truffles. Phylogenetic relationships and divergence time estimations obtained with 1,057 complete and single-copy BUSCO genes. The species tree was constructed using an amino acid supermatrix, while both protein (black) and nucleotide (red) data were used for divergence time estimation. All nodes received maximum support values from the species tree inference, except for the split between *Arthrobotrys* and Pezizomycetes. The outgroup *Schizosaccharomyces osmophilus* was removed for visualisation purposes.

Generally, our results are consistent with previous phylogenomics (Murat et al. 2018; Miyauchi et al. 2020; Shen et al. 2020) without over- or underestimation of divergence times of Ascomycota. Moreover, for most nodes we did not observe strikingly different results when using nucleotide or protein data, with all confidence intervals largely overlapping (Fig. 3).

Our estimates replaced the diversification of Tuberaceae (Tuber/Choiromyces split) toward the end of the Cretaceous (∼76 million years ago (MYA)), between previous estimations of ∼140 MYA (Bonito et al. 2013; Murat et al. 2018) and ∼45 MYA (Miyauchi et al. 2020). The dawn of diversification within the genus *Tuber* is the Paleogene, around ∼56 MYA, which sees the emergence of major subgeneric lineages, the Aestivum clade at ∼34 MYA and the Melanosporum clade at ∼26 MYA. The estimated split between *T. panzhihuanense* and *T. borchii* (Puberulum clade), its closest relative in the data set, is estimated to have occurred between ∼26 and ∼30 MYA.

### High and compositionally variable TE content in Tuberaceae

We estimated the TE content of the 13 included Pezizales species using automatically *de novo* generated libraries. Although we did not manually curate these elements, as previously shown, the overall TE content and its composition did not change significantly when using the raw and partially curated repeat library in the Chinese white truffle genome (Supplementary Figure S2), making our results suitable for comparing the TE content between different species.

Firstly, we explored the presence of possible biases in assembly sizes (as a proxy for genome size) and TE estimations due to different levels of assembly contiguities measured in terms of N50 values, and we did not identify any significant relationship (all *P* > 0.05, supplementary table S5). On the other hand, we found a strong positive correlation between assembly size and total TE content, as well as Gypsy content, also when correcting for shared evolutionary histories using phylogenetic independent contrasts (PIC; Fig. 4A-B, supplementary table S6; supplementary table S7). In particular, Tuberaceae genomes are characterised by higher TE content, dominated by Gypsy and LINE retrotransposons as well as by larger genomes compared to other analysed species (Fig. 4C; supplementary table S6). For other TE groups beside Gypsy, we found a positive and significant correlation with genome size only before PIC correction (supplementary table S7). Moreover, for total TE, DNA, and non-Gypsy LTR content, we found strong and significant phylogenetic signals with Pagel’s λ values approaching 1, whereas we did not find evidence of phylogenetic structure for Gypsy and LINE content (supplementary table S7). Indeed, closely related *Tuber* species can have drastically different contributions of Gypsy elements to the overall repeat content, even between recently diverging species, such as *T. melanosporum* and *T. indicum* (Fig. 4C). Similarly, while we identified members of all Gypsy clades also within Morchellaceae, *Gyromitra esculenta* Pers. ex Fr. and *Pyronema confluens* Tul. & C. Tul. (supplementary figure S11; supplementary table S6), their relative contribution to the total Gypsy content was highly variable across truffle species (Fig. 4D). Notably, the 4 identified clades cover the majority of the Gypsy content in Tuberaceae with a much greater contribution of CHD-related LTRs (Fig. 4D; supplementary table S6). These results suggest independent Gypsy expansions in different truffle lineages concurrently with their diversification. Lineage-specific activity of Gypsy clades in *T. panzhihuanense* was confirmed by repeat landscape profiles using a neutral substitution rate of 3.36 × 10E-3 (Fig. 4E). Here, we found that most of the Gypsy content is due to past but lineage specific activity with the high genomic occurrence of the Prometheus clade that mainly derives from one burst that occurred after the split with *T. borchii* (Fig. 3). We did not produce repeat landscape profiles for other species due to the significantly lower quality of their assemblies; however an evolutionary scenario mainly involving lineage-specific amplifications was supported by Gypsy phylogenetic analyses with the presence of highly dense species-specific clades that are evolutionarily distant between each other (Fig. 4F).

**Fig. 4.**
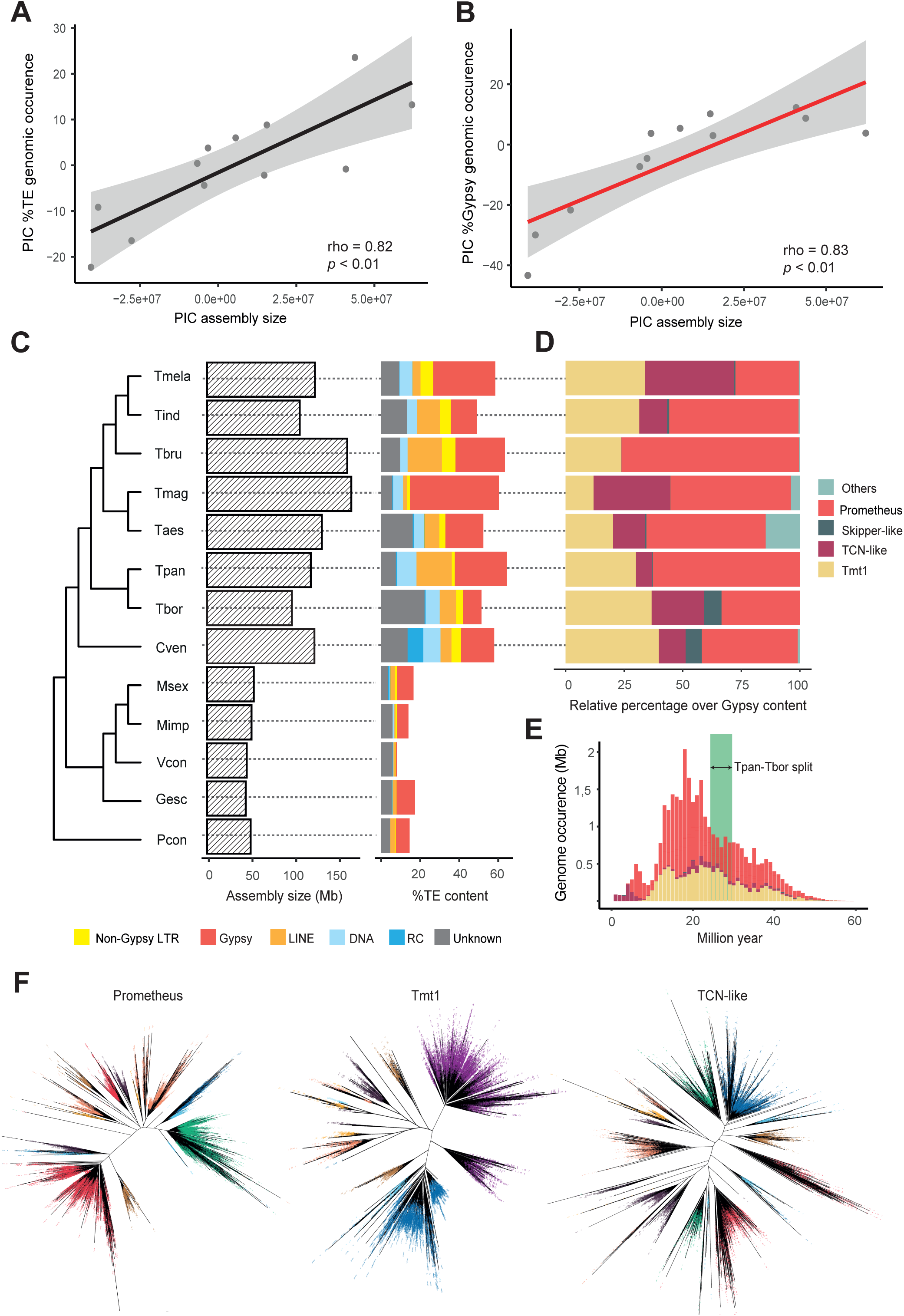
Lineage-specific proliferation of different Gypsy clades drives genome expansion in Tuberaceae. Correlations between **(A)** total TE content and **(B)** Gypsy content with assembly size, used as a proxy for genome size after correcting for shared evolutionary history using phylogenetic independent contrasts (PIC). **(C)** Assembly size and TE content for 13 Pezizales species used in comparative genomic analyses. Species abbreviations are reported in Fig. 3. **(D)** Relative contribution of identified clades to the total Gypsy content of Tuberaceae. Others referred to Gypsy clades that were not identified and characterised in *T. panzhihuanense*. **(E)** Repeat landscape profile describing the activity over time of Gypsy clades in the *T. panzhihuanense* genome after transforming the CpG-corrected Kimura distance into millions of years using a substitution rate of 3.36 × 10E-03. The green box highlights the mean divergence time between *T. panzihuhanense* from *T. borchii* based on nucleotides (lower bound) and amino acids (upper bound). **(F)** Clade-specific phylogenetic trees of all Gypsy RT segments with a length of at least 100 amino acids extracted from all Tuberaceae genomes. Different colours highlight different species.

### Synteny conservation between true truffles and Morchellaceae despite high gene-family turnover rates and TE accumulation

Murat et al. (2018) found a high rate of gene family turnover within Tuberaceae together with an unexpected high level of microsynteny conservation within them. Here, we took advantage of the increased number of truffle genomes together with our high quality *T. panzhihuanense* assembly to further test these hypotheses, their relationship with TE accumulation and study syntenic relationships at larger taxonomic scales, between true truffles and Morchellaceae. Because no gene annotation was publicly available for *Verpa conica* (O.F. Müll.) Sw and *G. esculenta* we performed a *de novo* annotation following the same pipeline used for *T. panzhihuanense* (See supplementary table S8 for summary statistics). We estimated 2 different rates of gene family turnover (CAFE λ) based on Orthofinder inferred orthogroups across Pezizales time tree, confirming a 4-fold higher turnover rate (0.163 vs 0.036) in Tuberaceae compared to other analysed species (Fig. 5A). Early evolution of Tuberaceae appeared to have been mainly impacted by gene losses, however, we also identified 27 significantly expanded gene families in their stem branch (CAFE Viterbi *p* < 0.05; Fig. 5A).

**Fig. 5.**
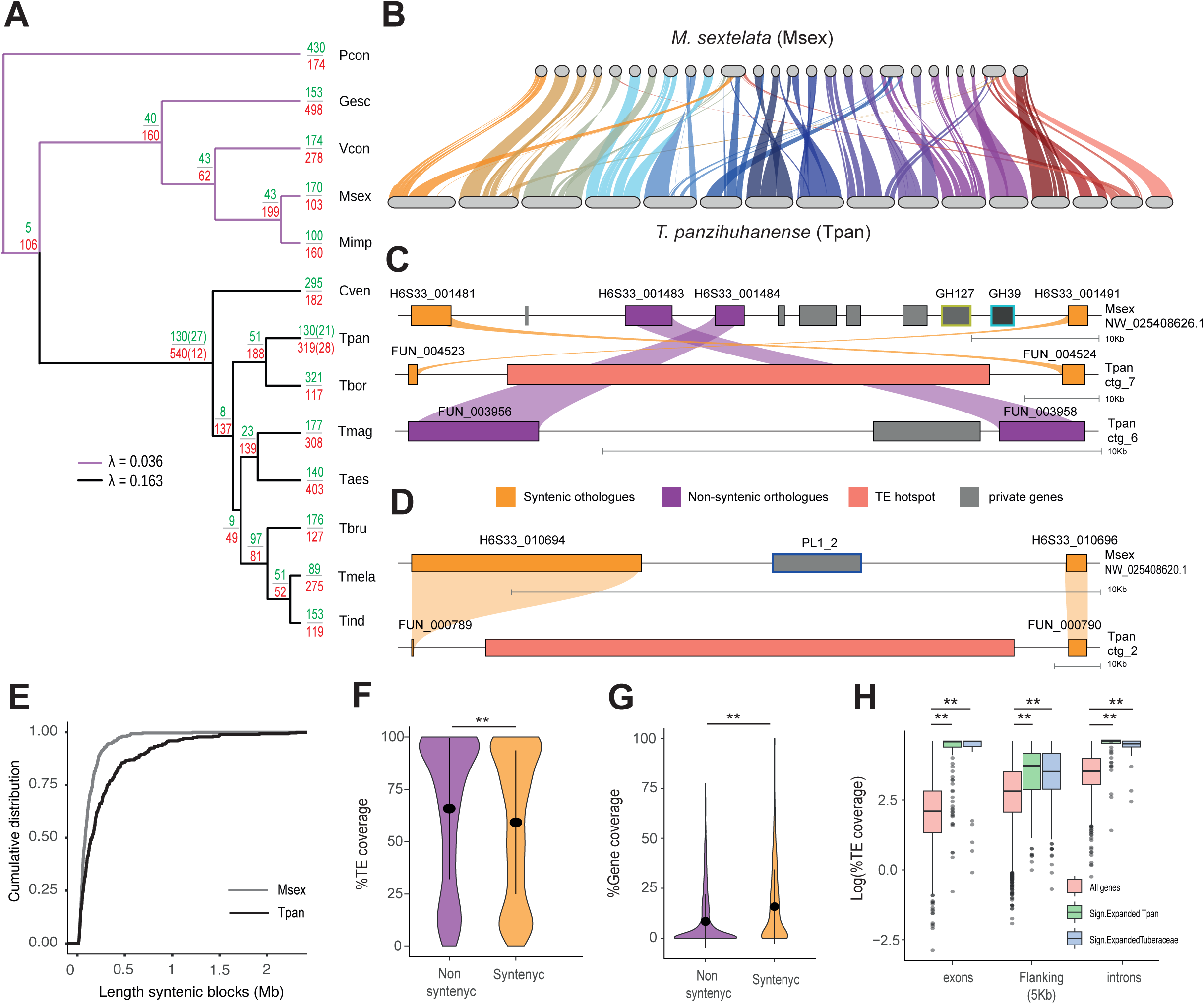
High gene family turnover rates within Tuberaceae and synteny conservation with Morchellaceae. **(A)** Gene family expansions and contractions along the Pezizales phylogeny. At each node, the top number indicates the number of gene family expansions, while the bottom number indicates the number of contractions. The numbers in parentheses represent the significantly expanded and contracted gene families. λ = birth-death model rates fitted with CAFE for Tuberaceae and all other Pezizales species. Species abbreviations are shown in Fig. 3. **(B)** Syntenic relationships between the *T. panzhihuanense* and *M. sextelata* genomes. The length of syntenic blocks is proportional to their actual length in base pairs. **(C)** and **(D)** Examples of syntenic genomic regions between *T. panzhihuanense* and *M. sextelata* involved in gene loss and small-scale rearrangements. Private genes = genes without any known orthologous relationship. Blocks have different scales and are shown at the bottom. **(E)** Cumulative distribution of the length of syntenic blocks in *T. panzhihuanense* and *M. sextelata*. **(F)** and **(G)** Comparison of TE and gene coverage (% of base pairs) across syntenic and non-syntenic genomic regions in *T. panzhihuanense*, calculated over non-overlapping genomic windows of 50 kb. Non-syntenic genomic regions were found to be significantly more TE-rich and less gene dense (Wilcoxon rank-sum test, *p* < 0.01). **(H)** TE coverage (% of base pairs) across exons, introns, and 5 kb of gene-flanking regions for genes belonging to significantly expanded gene families identified in the branch leading to Tuberaceae diversification, as well as in the *T. panzhihuanense* terminal branch, compared to all other genes. Genes belonging to significantly expanded gene families were found to be significantly more TE-rich across all genomic compartments (Wilcoxon rank-sum test, *p* < 0.01).

Gene-based synteny analysis between *T. panzhihuanense* and the long-read based high quality *Morchella sextelata* M. Kuo assembly (contig N50=1.8Mb) revealed relatively high levels of synteny between the 2 species (Fig. 5B; supplementary figure S12). Indeed, 61% of the *T. panzhihuanense* genome and 70% of single copy orthologs resulted syntenic with *M. sextelata* with syntenic block that can reach up to 2 Mb of length (mean = 269.5 kb; supplementary figure S13 A-B). Non syntenic genomic regions resulted 1.14 times more likely to contain TE hotspots according to chi-square statistics (sample size = 118,923,947; degree of freedom = 1, *X*^2^ = 1,632,590.5158 , *p* < 0.01), however, 59% of the TE hotspots are completely contained within syntenic blocks and flanked by syntenic genes (Fig. 5C-D). Visual inspection of some of these regions revealed some notable cases of loss of CAZy encoding genes in the *T. panzhihuanense* lineage related to small scale rearrangements (e.g. GH37 and GH39; Fig. 5C) and TE accumulation (e.g. PL1_2; Fig. 5D). Syntenic blocks are significantly longer in *T. panzhihuanense* compared to *M. sextelata* (Fig. 5E; Wilcoxon rank-sum test, *p* < 0.01) reaching a maximum value of 20 times increased length. Accumulation of TEs in intergenic genomic regions came out as the primary contributor to the increased genome size in Tuberaceae. Indeed, *M. sextelata* introns resulted slightly but significantly longer and more numerous compared to *T. panzhihuanense* (Wilcoxon rank-sum test, *p* < 0.01; supplementary figure S14 A-B), whereas intergenic genomic regions are on average ∼4.8-fold longer in the latter species (Wilcoxon rank-sum test, *p* < 0.01; supplementary figure S14 C), with a much stronger impact of TE accumulation on their length compared to introns (supplementary figure S14 D-E). Coherently with previous observations, syntenic genomic regions are slightly but significantly depleted in TEs (TE density: 0.59 vs 0.65; Wilcoxon rank-sum test, *p* < 0.01; (Fig. 5F) and almost two times more gene-dense compared to non-syntenic ones (Gene density: 0.159 vs 0.08; Wilcoxon rank-sum test, *p* < 0.01; Fig. 5G).

Finally, we found that genes belonging to significantly expanded families in the branch leading to Tuberaceae diversification as well as in *T. panzhihuanense* terminal branch, are significantly more TE-rich (Wilcoxon rank-sum test, all *p* < 0.01), in their exons, introns, and within 5 kb of flanking sequences (Fig. 5H). Additionally, genes included in TE-hotspots are enriched in paralogue copies of these fast-evolving families (chi-squared test, sample size = 8701, degree of freedom = 1, *X*^2^ = 149.1907, *p* < 0.01), with 4.7 times more of these genes than expected by chance. These findings led us to hypothesise that genes located within TE hotspots are likely to be members of multi-copy gene families. Indeed, 54% of the gene content of TE hotspots belongs to families with more than one gene, compared to 18% of the genome.

### Gene family expansions at the root of Tuberaceae are related to the establishment of an ectomycorrhizal lifestyle

Gene duplication can lead to neofunctionalization and protein diversification, potentially facilitating the emergence of novel phenotypic traits (Copley 2020). We therefore tested the hypothesis that gene duplication may have contributed to the transition from a saprotrophic to an ECM lifestyle in the Tuberaceae stem branch. To do this, we identified gene families associated with ECM (ECM-induced families) by intersecting our gene family set with genes found to be upregulated in *T. magnatum* ECM compared to free-living mycelium by Murat et al. (2018). We identified 328 ECM-induced gene families, of which 18 are expanded, and 5 of these are significantly expanded in the branch of interest (Fig. 6A; Supplementary Table S9; Supplementary Figure S15). Given that a total of 6,128 gene families are inferred by CAFE to be present at the root of the species tree and contain at least one Tuberaceae species (i.e., they are also present at the Tuberaceae root), we observed a significant overrepresentation of ECM-induced gene families by 2.5-fold and 3.7-fold among expanded and significantly expanded gene families, respectively (Fisher’s exact test; p < 0.01).

**Fig. 6.**
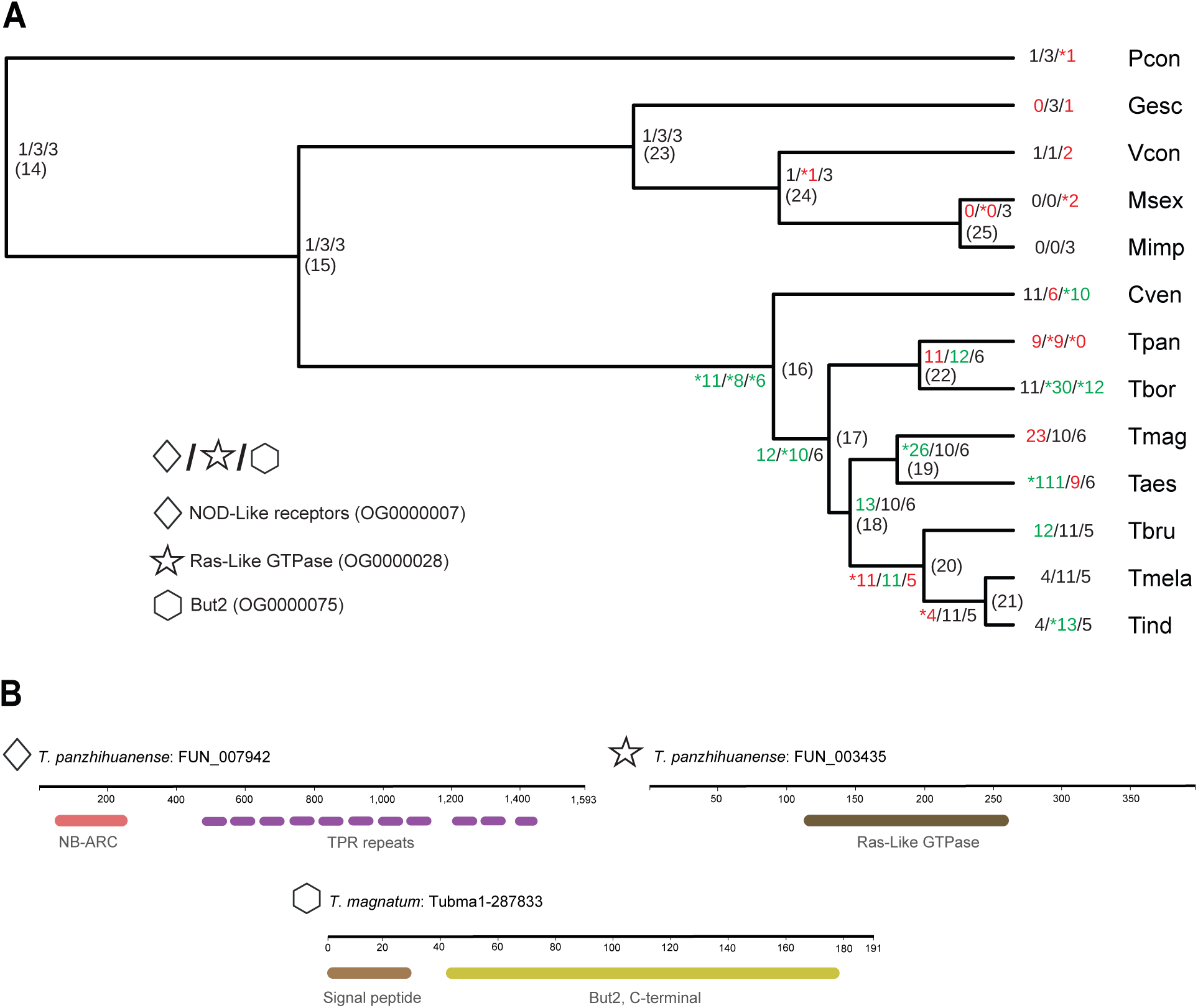
Ancestral gene duplications of ECM-induced gene families. **(A)** CAFE results for 3 ECM-induced gene families significantly expanded in the stem branch of Tuberaceae. For each node, the inferred and observed gene family counts are reported for internal and terminal branches, respectively. Green colours highlight expansion events, red colours indicate contractions, and significant changes are marked with an asterisk. In brackets is reported the node number and refers to supplementary table S9. Species abbreviations are reported in Fig. 3. **(B)** InterProScan domain annotation of representative proteins for each of the 3 gene families. The symbols correspond to the annotations in (A). In Supplementary figure S15 can be found the results for the other 2 significantly expanded ECM-induced gene families.

Rapid adaptation to a changing environment during ECM establishment has been suggested to involve signal transduction pathway cascades (Martin and Tunlid 2009). The significantly expanded gene family OG0000007 encodes NOD-like receptors (NLRs), which are a class of receptor proteins involved in various types of biotic interactions such as self/nonself recognition and ECM symbiosis (Dyrka et al. 2014). In *T. melanosporum* ECM, a high number of NB-ARC NOD-like receptors were also found to be upregulated (Martin et al. 2010). A total of 187 genes were clustered in this gene family, with one orthologue in *P. confluens* and one in *V. conica*. Morchellaceae have apparently lost this gene family, whereas Tuberaceae possess a variable but high number of paralogue copies. The genome of the ECM fungus *Laccaria bicolor* encodes a wide set of protein kinase and RAS small guanosine triphosphatase (GTPase) genes (Martin et al. 2008), with frequent neofunctionalization of paralogous copies and few pseudogenization events (Rajashekar et al. 2009). The expression of one of these genes, *Lbras*, depends on the interaction with host roots and it is expressed in established mycorrhizal tissues (Sundaram et al. 2001). We also found that Tuberaceae experienced an ancestral gene family expansion of putative Ras-like proteins (OG0000028). The gene family OG0000075 encodes for a wide set of But2-like proteins across Tuberaceae, however, *T, panzhihuanense* has apparently lost this gene family. But proteins of *Saccharomyces cerevisiae* are seemingly involved in the activation of the NEDD8 ligation pathway, which leads to substrate NEDDylation (Yashiroda and Tanaka 2003), a post-translational modification that is crucial for fungal cellular processes such as development and secondary metabolism (Yang et al. 2022). But2 has been poorly studied but it is known that many proteins show similarity with its C-terminal region (Yashiroda and Tanaka 2003). Murat et al. (2018) found one gene encoding for a SSP But2-like protein overexpressed in *T. magnatum* ECM which clustered within this gene family (TubMa1-287833). SP proteins with high similarity to But2 C-terminal domain could be involved, with other fungal secreted proteins, in affecting host transcription and responses needed for the establishment and retention of mycorrhizal structures.

## Discussion

Here, we present a high-quality genome assembly of the Chinese white truffle *T. panzhihuanense*, which represents the most contiguous genome of a Pezizales species assembled to date. Thanks to its high contiguity, our recalibrated divergence time, and in-depth analyses of TEs and gene family evolution across Pezizales, we could explore with unprecedented detail the impact of massive TE burst in genome architecture and evolution of Tuberaceae.

Based on multiple fossil calibration points and a genome-scale matrix, we placed the diversification of Tuberaceae (∼76 Mya) and true truffles (∼56 Mya) in a more biologically meaningful timeframe compared to previous studies (Bonito et al. 2013; Murat et al. 2018: Miyauchi et al. 2020) which were based on substitution rates inferred from distantly related taxa and/or few fossil data. A late Cretaceaous emergence of the family is indeed consistent with the angiosperm terrestrial revolution (Benton et al. 2021) that gave rise to the most species-rich living clades of angiosperms (Ramírez-Barahona et al. 2020), mammals (Álvarez-Carretero et al. 2022), birds (Jarvis et al. 2014) and arthropods (Benton et al. 2021; Montagna et al. 2019) but with their following diversification delayed until the Paleogene. The diversification of truffles took therefore place in a context of expanding angiosperm-based ecosystems in which these fungi coevolved and codiversified with their host plants and the animals responsible for their spore dispersal (Benton et al. 2021; Brundrett and Tedersoo 2018). It is noteworthy that within Basidiomycota, ECM Agaricomycetes apparently experienced a similar evolutionary course diversifying rapidly during Paleogene following a late Cretaceous divergence (Sato 2023).

Our re-analysis of 13 Pezizales genome assemblies confirms previously observed patterns (Martin et al. 2010; Murat et al. 2018; Miyauchi et al. 2020), finding a strong and significant correlation between TEs, particularly Gypsy LTR retrotransposons, and genome size. Consistent with previous studies on the Périgord black truffle *T. melanosporum* (Payen et al. 2016), and similarly to other fungi (Muszewska et al. 2011), chromoviridae-related clades have been particularly successful in expanding within truffle genomes. Here, we additionally found representative elements of all 4 identified clades across multiple Pezizales species, suggesting that the great majority of truffle LTRs derive from pre-existing lineages with CHD elements that underwent massive, lineage-specific proliferations concurrently with their diversification, possibly due to demographic histories (Murat et al. 2010) and stochastic activation of specific lineages (Oggenfuss et al. 2021).

Increasing TE content is widely associated with decreased gene density, with Tuberaceae genomes representing one of the most iconic cases within fungi (Muszewska et al. 2019). Analysing the Chinese white truffle assembly, we found that more than half of its genome is composed of long stretches of heterochromatic DNA, dominated by long LINE Tad1 and LTR Gypsy insertions. Non-random distribution of TEs, particularly Gypsy LTRs, was already observed in *T. melanosporum* (Martin et al. 2010; Payen et al. 2016). Indeed, chromoviridea-related LTRs have a strong preference for heterochromatic genomic regions in plants (Neumann et al. 2011) and in the fungus *Schizosaccharomyces pombe* **Lindner**, where the CHD domain directly recognizes histone H3 K9 methylation (Gao et al. 2008). No heterochromatic insertion preference has been described so far for LINE Tad1 elements, and their massive presence within these regions can be explained by negative selection against long insertions in close proximity to genes (Buckley et al. 2017; Ruggieri et al. 2022; Martelossi et al. 2024). Instead, host PCGs are clustered in genomic regions highly depleted in TEs (in particular long retrotransposon insertions), where gene and TEs densities reach similar levels to those in less TE-rich fungi. This process of retrotransposons-mediated self-perpetuating expansion of heterochromatic genomic regions (Gao et al. 2008), contrary to intron gain or intron expansion, is also the main driving force behind the increased genome size of truffles. Genome expansion is often counteracted by DNA loss following an "accordion" model of genome evolution (Kapusta et al. 2017). Here, similar to what was previously observed in the fungus *Pleurotus ostreatus* **(Jacq.) P. Kumm.** (Castanera et al. 2016), we found evidence of non-allelic homologous recombination between LTR regions of Gypsy transposons. It is therefore likely that Tuberaceae genomes coped with TE expansions both due to strong insertion preference of highly active Gypsy elements and, on the host genome side, due to MIP and an increased rate of ectopic recombination resulting from the spread of homologous sequences throughout the genome, limiting the deleterious effects of an uncontrolled proliferation.

The emergence of non-homologous and rearranged genomic regions is expected to be one of the main outcomes of TE accumulation (Thon et al. 2006; Aguileta et al. 2009; Treindl et al. 2021). Coherently with this expectation, non-syntenic genomic regions between *T. panzhihuanense* and *M. sextelata*, a Morchellaceae with low TE content and high gene density (Han et al. 2019), harbours a higher number of TEs and a lower number of genes compared to syntenic ones. On the other hand, most of the orthologous genes identified between the two species, as well as most of their genomes, are syntenic despite the aforementioned TE accumulation and more than 200 million years of divergence. Indeed, TE accumulation resulted in a drastically increased length of syntenic blocks, but a conserved mid-scale synteny, with the occasional emergence of small-scale genomic rearrangements, possibly favouring the establishment of novel TE-rich heterochromatic DNA. The GH127, GH39, and PL1_2 CAZy enzymes are encoded by *M. sextelata* but have been lost in *T. panzhihuanense* suggesting that TE insertions, together with sequence decay (Murat et al. 2018) might have contributed to gene loss in the clade. Despite the absence of chromosome-scale assemblies to dissect intra- and inter-chromosomal genomic rearrangements, our results recapitulate the mesosynteny pattern formally described in filamentous fungi (Hane et al. 2011) and also observed between *T. melanospurum* and the Eurotiomycetes *Coccidioides immitis* **Rixford & Gilchrist**, (Martin et al. 2010). This mode of chromosome evolution is expected to occur when intra-chromosomal recombination events, such as inversions and gene loss, are more frequent than inter-chromosomal recombination events, such as translocations (Hane et al. 2011). On the other hand, we also found that multi-copy gene families are preferentially located within TE-rich genomic regions, reinforcing the idea that they might act as hotspots for copy number variation (Montanini et al. 2014). Interestingly, the overrepresentation of ECM-induced gene families among the significantly expanded ones in the stem branch of the family suggests that protein diversification due to neo and/or subfunctionalization may have contributed to the establishment of an ECM lifestyle. Overall, these findings demonstrate the primary role of TEs in the high gene-family turnover rate observed in Tuberaceae and raise the necessity of studying co-expression profiles of genes and nearby TEs in future studies on the establishment and maintenance of Tuberaceae ECM.

Finally, we explored the structure of nuclear rDNA loci, which were difficult to characterise in previous genome assemblies due to the use of only short-read data. Similar to most, but not all, other Pezizomycotina (Bergeron and Drouin 2008), truffles exhibit the 5S ribosomal gene dispersed throughout the genome and not included in the 45S rDNA array. The high level of similarity found between 18s, 5.8S and 28S copies, but not in the 5S paralogues, and the presence of partial arrays only at the ends or within rDNA-only contigs, suggest that these genes are undergoing concerted evolution due to high homogenization within a single cluster (Ganley and Kobayashi 2011; Hori et al. 2021). The complex IGS region, similarly to what is observed in humans and budding yeast (Ganley and Kobayashi 2011; Hori et al. 2021), is composed of tandemly arranged repeats and can undergo deletion and duplication events, leading to rDNA arrays of different length. We also provide evidence of intragenomic variability of 45S rDNA units, with the identification of one diverging copy that is likely located outside the presumptive main cluster. Its existence should not invalidate the use of the ITS region for species and strain identification in *T. panzhihuanense* using classic Sanger sequencing as, even if we assume a successful primer binding, most of the fluorescent intensity of the electropherogram would come from the paralogous copies of the main clusters. However, it can become a problem with metagenomics studies based on environmental DNA samples (Bradshaw et al. 2023). We therefore encourage future investigation of the 45S rDNA gene polymorphisms in truffles to avoid the establishment of new cryptic species and invalid diversity estimates due to intra-genomic polymorphisms.

## Material and Methods

### Sampling and DNA/RNA sequencing

The sample used for genome sequences was collected in 2020 by the Jinsha river basin (river section of Zhongba village, Renhe district, city of Panzhihua, China). The fruiting body was isolated and taxonomically identified based on morphology and molecular characteristics and sent to Personalbio Co., Ltd. (Shanghai, China) for whole Genome Sequencing (WGS). WGS was performed both on a PacBio Sequel II and an Illumina NovaSeq 6000 platform generating subreads and obtaining 38+ Gb of HiFi reads. RNA-Seq data used for genome annotation were obtained from *T. panzhihuanense* ectomycorrhizal synthesis with *Pinus massoniana*. RNA was extracted from 3 replicates of each 100 individual ECMs isolated from one *P. massoniana*-*T. panzhihuanense* mycorrhizal seedlings. With samples sent to Novogene Technology Co., Ltd. (Beijing, China), the library constructed was sequenced on Illumina NovaSeq 6000 platform in 150 PE mode. For detailed sample identification, mycorrhizal synthesis preparation, and library construction operations, see the supplementary material: Supplementary Material and Methods for detailed explanations.

### Genome survey and assembly

Illumina reads were quality checked with fastQC v0.11.7 (Andrews 2010), and trimmed with Trimmomatic v0.39 (Bolger et al. 2014) (*-phred33 LEADING:3 TRAILING:3 SLIDINGWINDOW:4:15 MINLEN:36*). Trimmed reads were used with KAT v2.4.2 (Mapleson et al. 2017) *hist* subtool to produce k-mer frequency histogram. Genome size and heterozygosity were estimated with GenomeScope 2.0 (Ranallo-Benavidez et al. 2020). PacBio subreads were processed to obtain final CCS (Circular Consensus Sequences) sequencing using PacBio CCS tool v6.4.0 (*--min-rq 0.99*). The genome was assembled with Hifiasm v0.16.1 (Cheng et al. 2021) under default parameters and Blobtools v1.1 (Laetsch and Blaxter 2017) was used to identify contaminant contigs. To obtain coverage information, we mapped HiFi reads back to the assembly with Minimap2 v2.24-r1122 (Li 2018) (*-ax map-hifi*), while for taxonomic annotation, we blasted all contigs against the NCBI nt database (*Blastn -max_target_seqs 10 -max_hsps 1 -evalue 1e-25*). Reads mapped to contigs identified as contaminants were excluded and the genome reassembled with Hifiasm. Quality and completeness of the final assembly were assessed with BUSCO v5.2.2 (Mosè et al. 2021; fungi_odb10 reference database) and Merqury (Rhie et al. 2020) with a k-mer size of 21.

### Repeat annotation

For repeat annotation, we produced a starting raw repeat library with RepeatModeler2 v2.0.4 (Flynn et al. 2020) with the LTR structure extension. Raw consensus sequences were used for a first genome annotation with RepeatMasker v4.1.2-p1 (Smit 2013-2015) in sensitive mode (*-s*). Based on RepeatMasker results, we isolated all consensus sequences with at least 50 instances in the genome and/or characterised by protein fragments based on Blastx (E-value 1E-05) results against the RepeatPeps database from the RepeatMasker package. Selected sequences were subjected to manual curation and classification at the class level following a “Blast-Extend-Extract” process as described in Goubert et al. (2022) and Peona et al. (2024). Curated and uncurated consensus were merged and redundancy removed following the 80-80 rule (i.e. requiring a minimum 80% identity along the 80% of the shortest sequence; Wicker et al. 2007; Goubert et al. 2022) with cd-hit-est (*-c 0.8 -n 5 -aS 0.8 -g 1 -G 0 -t 1*). The merged library was used for repeat annotation with RepeatMasker in sensitive mode. We post-processed RepeatMasker results with the parseRM.pl script (https://github.com/4ureliek/Parsing-RepeatMasker-Outputs/blob/master/parseRM.pl) to obtain genome-wide repeat estimations and RepeatCraft (Wong and Simakov 2019) in *loose* mode to defragment closely spaced repeat loci and obtain a final TE annotation.

### Gene annotation

Quality of RNA-Seq reads was assessed with fastQC v0.11.7 and trimmed with Trimmomatic v0.39 (-phred33 LEADING:3 TRAILING:3 SLIDINGWINDOW:4:15 MINLEN:36). We assembled the transcriptome with TRINITY v2.1.1 (Grabherr et al. 2011) under default parameters.

For gene annotation, we used Funannotate v1.8.16 (Palmer and Stajich 2019) performing an *ab-initio* gene prediction with Augustus (Stanke et al. 2008), SNAP (Korf 2004), GlimmerHMM (Majoros et al. 2004), CodingQuarry (Testa et al. 2015) and GeneMark-ES (Lomsadze et al. 2005). The assembled transcriptome, 4 Ascomycota RefSeq proteomes available on NCBI (GCF_000151645.1: *Tuber melanosporum*, GCF_003444635.1: *Morchella importuna*, GCF_020137385.1: *Morchella sextelata*, and GCF_024521635.1: *Tricharina praecox*) and the Swiss-Prot database (The Uniprot Consortium 2023) were used as external evidence to improve the annotation process.

Functional annotation of predicted genes was performed with InterProScan V5.70 (Jones et al. 2014) to obtain PFAM domains (Mistry et al. 2021) and InterPro entries. CAZymes were separately annotated with dbCAN3 (Zheng et al. 2023). rDNA genes were annotated using RNAmmer (Lagesen et al. 2007). Secreted proteins (SP) were annotated following the method of Pellegrin et al. (2015). We considered SP proteins as small secreted proteins (SPPs) when their length was equal to or lower than 300 aa, as in Pellegrin et al. (2015).

### TE genomic distribution analyses

To test the interplay between gene density and TE densities we subdivided the *T. panzhihuanense* nuclear genome into non-overlapping windows of 50 Kb using bedtools *makewindows* (Quinlan and Hall 2010) and calculated the proportion of base pairs occupied by genes and transposons for each interval.

To detect genomic regions significantly enriched (TE hotspots) and depleted (TE coldposts) in TEs we applied a binomial test on the number of base pairs occupied by transposons in genomic sliding windows of 20 kb with a window step size of 5 kb. For each window, we computed the TE coverage using bedtools *coverage* and compared the observed number with the genome-wide average estimation computed across all windows. For each genomic interval, we separately tested for both significant greater and lower TE content compared to genome-wide estimation as alternative hypotheses, generating two sets of intervals representing genomic regions significantly enriched and depleted in TEs, respectively. *P*-values were corrected for multiple testing with the Benjamini–Hochberg procedure and a false discovery rate (FDR) cut-off of 0.01. Because applying a sliding window approach may result in overlapping genomic intervals being annotated as both TE-rich (TE hotspots) and TE-depleted (TE coldspots), we used bedtools *subtract* reciprocally on both interval sets to remove overlapping regions. Genes completely falling within TE-enriched or TE-depleted windows were extracted using bedtools *intersect* (-f 1).

### Identification, classification and genome annotation of LTR Gypsy families

We reconstructed *T. panzhihuanense* Gypsy families based on the evolutionary relationship of RT domains of single insertions. We isolated RT nucleotides and protein segments longer than 400 nt through homology searches (BLASTx; E-value 1E-05) querying all LTR insertions against a database of RT domains downloaded from GypsyDB (Llorens et al. 2011). Gypsy RTs were further confirmed blasting back all sequences (Blastp E-value 1E-05) against the RepeatPep library and keeping only queries with a best-hit against a Gypsy transposon.

For phylogenetic analyses, we reduced the size of the dataset by clustering all confirmed Gypsy RT protein sequences with cd-hit at an identity threshold of 80% (*-c* 0.8). Representative sequences were aligned with MAFFT v7.520 in G-INS-i mode (Katoh and Standley 2013) and columns with more than 20% gaps were excluded with trimAl v1.4.rev22 (*-gt* 0.8; Capella-Gutiérrez et al. 2009). A maximum likelihood tree was inferred with IQ-TREE v2.2.2.6 (Minh et al. 2020) using ModelFinder (Kalyaanamoorthy et al. 2017) to identify the best-fit evolutionary model and performing 1,000 ultrafast bootstrap replicates to assess nodal support (Hoang et al. 2018).

To complement the phylogenetic analyses, we built a network based on pairwise alignment relationships from an all-to-all Blastn (E-value 1E-06) analysis of the nucleotide sequences of the RT segments. Alignments shorter than 300 bp were excluded, and bit scores were used as edge weights to find the community partition with the largest modularity, as implemented in the *greedy_modularity_communities* function of the NetworkX Python package (Hagberg et al. 2008). Gypsy communities were mapped to the RT phylogenetic tree and we manually defined Gypsy families as highly divergent and supported clades (Bootstrap ≥ 75), eventually splitting previously identified communities.

To build consensus sequences representative of identified families, we recovered all insertions from which we derived the RT segments included in the phylogenetic analyses and separately aligned them with MAFFT v7.520. From these alignments, we built starting consensus sequences with CIAlign (Tumescheit et al. 2022). We then refined these consensus with a “Blast-Extend-Extract” process, as previously described, and we finally checked for their completeness with TE-aid (Goubert et al. 2022).

We classified curated Gypsy families relying on phylogenetic relationships with previously described elements. Briefly, RT protein domains extracted from each consensus were added to a set of reference Gypsy elements including known Gypsy clades described in GypsyDB and elements from Novikova et al. (2010) and Riccioni et al. (2008). Sequences were aligned with MAFFT G-INS-i and gappy positions removed with trimAl (*-gt* 0.8). We inferred a ML tree with IQ-TREE together with ModelFinder and 1,000 UltraFastBootstrap replicates. Each Gypsy family was assigned to the closest known clade if the bootstrap value was ≥ 75.

Classified Gypsy families were used in an additional RepeatMasker analysis in sensitive mode and results were post-processed to obtain their genome-wide estimations and repeat landscapes, as previously described. We used the parseRM.pl script to produce a repeat landscape describing the activity of transposons through absolute time using a Tuber-specific neutral substitution rates (See Material and Methods: Phylogenomics and divergence time estimation)

### Identification of autonomous Gypsy copies and estimation of NAHR events

We screened for putatively autonomous Gypsy transposons based on the repeat annotation obtained with the consensus sequences of classified Gypsy families. We blasted back (BLASTn, E-value 1E-05) all insertions against their source consensus and considered an element as a full length when the alignment reciprocally covers the insertion and its consensus sequence for 90% of their length. Putative autonomous elements were identified by firstly self-blasting full length elements to identify LTRs regions. For those with confirmed LTRs, ORFfinder (https://www.ncbi.nlm.nih.gov/orffinder/) was used to extract translated open reading frames (ORFs) longer than 300 aa. RT, Integrase (INT) and Ribonuclease (H) domains were identified via hmmscan v3.4 (E-value 1E-05; http://hmmer.org/) against their HMM profiles downloaded from GypsyDB. We considered as putative autonomous elements those encoding for RT, INT and RNase H domains within a single ORF. We classified an insertion as a solo-LTR arising from an NAHR event when the alignment reciprocally covers both the insertion and the LTR region of the source consensus for at least 90% of their length. All other instances were considered as degenerated and/or fragmented elements.

## Comparative genomics analyses

### Phylogenomics and divergence time estimation

We performed species tree inference across a dataset of 31 Ascomycetes genomes available on NCBI. Species were chosen in order to have as many calibration points as possible for the divergence time estimation (supplementary table S11).

We extracted single-copy orthologous genes from all 31 fungal genomes using BUSCO and the Ascomycota_odb10 reference database. Amino acid and nucleotide sequences of complete and single-copy BUSCO genes present in at least 95% of the species were aligned using MAFFT in auto mode and cleaned of gappy positions with trimAl (-automated1). Single-gene amino acid alignments were then concatenated and subjected to maximum likelihood (ML) tree inference with IQ-TREE, incorporating ModelFinder2 and 1,000 UltraFast Bootstrap replicates.

To obtain a reliable timeframe for Ascomycota and, specifically, truffle diversification, we applied a Bayesian approach using multiple fossil calibrations as priors on our genome-scale dataset and the previously inferred species tree, as implemented in the MCMCTree package (dos Reis and Yang 2019). Specifically, we used 10 fossils with unambiguous phylogenetic placements, fitting a flat distribution between minimum and maximum bounds, and a normal distribution for the root of the tree (split between Taphrinomycotina and Saccharomycotina + Pezizomycotina). All priors are detailed in supplementary table S11, along with a deep justification for each fossil in supplementary material: Supplementary Material and Methods. Three runs were performed under an independent rates model, and convergence of the MCMC chains was assessed using Tracer (Rambaut et al. 2018). MCMCTree was run on both amino acid and nucleotide alignments to assess the sensitivity of the analyses to the underlying dataset.

To estimate a putative neutral substitution rate for Tuberaceae, we extracted all 4-fold degenerate sites from the previously identified Tuberaceae BUSCO genes and concatenated them. LSD2 (To et al. 2016) within IQ-TREE2 was then used to estimate a fixed substitution rate within the Tuberaceae subtree, calibrating all nodes setting as maximum and minimum bound the 95% highest posterior density estimation obtained from MCMCTree analyses on the nucleotide alignment.

### Repeat annotation and analyses across additional Pezizales genomes

All Pezizales genomes included in phylogenetic analyses were used in a *de novo* repeat discovery with RepeatModeler2 and the LTR structure extension. Raw species-specific repeat libraries were used for repeat annotation in each genome with RepeatMasker to obtain a rough estimation of their repetitive content. Relationships between genome size, and TE content were assessed with correlation analyses to the raw and to the data corrected for shared evolutionary histories using phylogenetic independent contrasts (PIC) with the *pic* function of Ape R (R core team 2021) package (Paradis et al. 2004). Phylogenetic signal was tested with Pagel’s λ (Pagel 1999) with the *phylosig* function of the Phytools R package (Revell 2024).

To characterise the distribution of different Gypsy clades across Pezizales diversity, we classified all Gypsy consensus sequences following the same approach used to classify Gypsy families isolated from the *T. panzhihuanense* genome after adding the *T. panzhihuanense* classified Gypsy families to the reference RT database.

Classified consensus sequences mined from Tuberacaeae species were subjected to manual curation following a “Blast-Extend-Extract” process, as previously described. Curated consensus were used in an additional RepeatMasker analysis on their source genome to obtain an accurate estimation of their genomic occurrence. RT fragments longer than 400 nt were extracted from the annotation (BLASTx; E-value 1E-05) and elements coming from Prometheus, Tmt1 and TCN-like clades were separately aligned via MAFFT in G-INS-i mode and, after removed gappy positions with trimAl (*-gt* 0.8), subjected to phylogenetic inference with FastTree v2.1.10 (Price et al. 2010) under a GTR + Gamma model.

### Gene family evolutionary analyses and syntenic genomic region detection

Syntenic genomic regions between *T. panzhihuanense* and *M. sextelata (*GCF_024521635.1) genomes were identified with GENESPACE v1.3.1 under default parameters (Lovell et al. 2022) after excluding the identified ribosomal contigs of *T. panzhihuanense*. A riparian plot was generated from inferred syntenic blocks with the *plot_riparian* function with the length of the block proportional to their actual length in base pairs. Syntenic and non-syntenic orthologs were extracted from GENESPACE results with the *query_pangenes* function.

We studied gene families turnover dynamics during Pezizales diversification using CAFE5 (Mendes et al. 2020) and the corresponding nucleotide divergence time estimation as background tree. Gene family counts were obtained inferring orthogroups from the proteomes of all species with Orthofinder v2.5.5 (Emms and Kelly 2019) with diamond (Buchfink et al. 2015) in ultra sensitive mode (*-S diamcond_ultra_sens*). Because no gene annotation was freely available for *Verpa conica* (GCA_033030425.1) and *Gyromitra esculenta* (GCA_038503075.1), we performed a *de novo* annotation using Funannotate following the previously described procedure for *T. panzhihuanense*. Briefly, RNA-Seq reads were downloaded from NCBI (SRR12605086 and SRR5491178 for *V. conica* and *G. esculenta* respectively) and the transcriptomes were assembled with TRINITY v2.1.1 (default mode). Assembled transcripts and previously described protein evidence (See Material and Methods: “Gene annotation”) were then supplied to Funannotate as external evidence. Gene families turnover analyses were performed estimating an error model to account for non-biological factors in gene families counts and specifying two separate lambdas (λ) for the tree, one for Tuberaceae and one for other Pezizales branches.

### Spatial relationships between TEs, syntenic genomic regions, and fast-evolving gene families

To explore the spatial relationships between TEs, synteny, and gene families evolution we looked at the genomic occurrence of transposons based on the RepeatMasker analyses obtained with the curated TE library on the *T. panzhihuanense* genome. After extracting syntenic and non-syntenic genomic regions between *T. panzhihuanense and M. sextelata* from GENESPACE results, we compared their gene and TE density across non-overlapping genomic windows of 50 kb. Accumulation of TEs in significantly expanded/contracted gene families was tested by looking at the percentage of base pairs annotated as TEs across exons, introns and 5 kb flanking regions compared to the rest of the genes.

## Data availability

The *T. panzhihuanense* genome assembly, all gene annotations, as well the transposable element libraries and orthogroup clusters described in this article are available on Figshare at https://figshare.com/s/a435a5ac8371bcea2cae. The *T. panzhihuanense* genome assembly is available together with all generated reads in the NCBI database and can be accessed within the Bioproject PRJNA1110184.

## Acknowledgements

The authors would like to thank China Scholarship Council-University of Bologna Cooperation Scholarship. Part of the research was completed thanks to the Master of Science course in Microbiology, China, with the relevant degree thesis [Biological Traits of Occurrence of Chinese Truffles (*Tuber* spp.) in the Natural Truffle Producing Area of Jinsha River Basin, China] confidential until June 2025 at https://www.cnki.net/index/. A.T. has contributed to the present work during and with the support of the Italian national inter-university PhD course in Sustainable Development and Climate change (link: www.phd-sdc.it).

## Fundings

The topic was selected as one of the 2016-2017 Sichuan Agricultural University Subject Construction Dual Support Projects (Special funding projects for outstanding young researchers) led by Y.H. and X.Z. This work was partially supported by the Canziani request funded to F.G. and the ‘Ricerca Fondamentale Orientata’ (RFO) funding from the University of Bologna to F.G. A.T. contributed to the present publication while attending the PhD programme in Sustainable Development And Climate Change at the University School for Advanced Studies IUSS Pavia, Cycle XXXVIII, with the support of a scholarship financed by the Ministerial Decree no. 351 of 9th April 2022, based on the NRRP - funded by the European Union - NextGenerationEU - Mission 4 "Education and Research", Component 1 "Enhancement of the offer of educational services: from nurseries to universities” – Investment 4.1 “Extension of the number of research doctorates and innovative doctorates for public administration and cultural heritage”.

## Author contributions

Y.H., X.Z., J.M., J.V., F.G., and A.Z. designed the study. Y.H., K.X., Y.C., and X.Z. sampled the specimens and prepared the materials used for genomic and transcriptomic extraction. Y.H., K.X., Y.C. performed RNA extraction; J.V. performed genome assembly and gene annotations; J.M. performed transposable elements and comparative genomic analyses. A.T. and O.R.S. selected the species and the fossils for divergence time estimation. J.M. and A.T. performed divergence time estimation. O.R.S. and F.G. supervised the analyses. J.M., J.V. Y.H. and A.T. prepared the figures and supplementary materials. J.M., J.V., A.T., Y.H., F.P., O.R.S., F.G., and A.Z. interpreted the results. J.V., J.M., and Y.H. wrote the first version of the manuscript. All authors critically revised the manuscript.

## Notes

https://figshare.com/s/a435a5ac8371bcea2cae

### Competing Interest Statement

The authors have declared no competing interest.

