## Supplementary Figures for "The high quality Chinese white truffle genome and novel fossil-calibrated estimate of Pezizomycetes divergence reveal the tempo and mode of true truffles genome evolution"

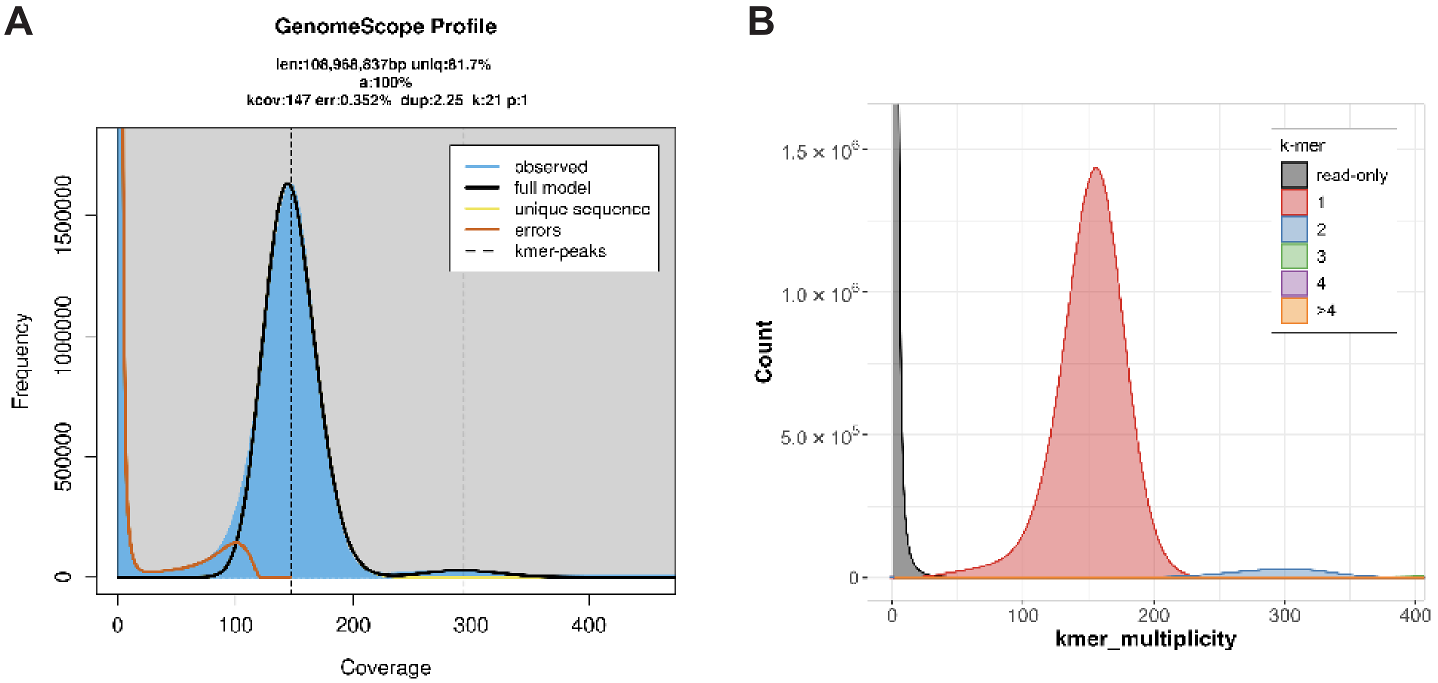


**Supplementary Figure S1**: (**A**) Short-reads kmer-based genome survey using genomescope with a kmer size of 21. (**B**) Copy number kmer spectrum obtained with Merqury and a kmer size of 21.

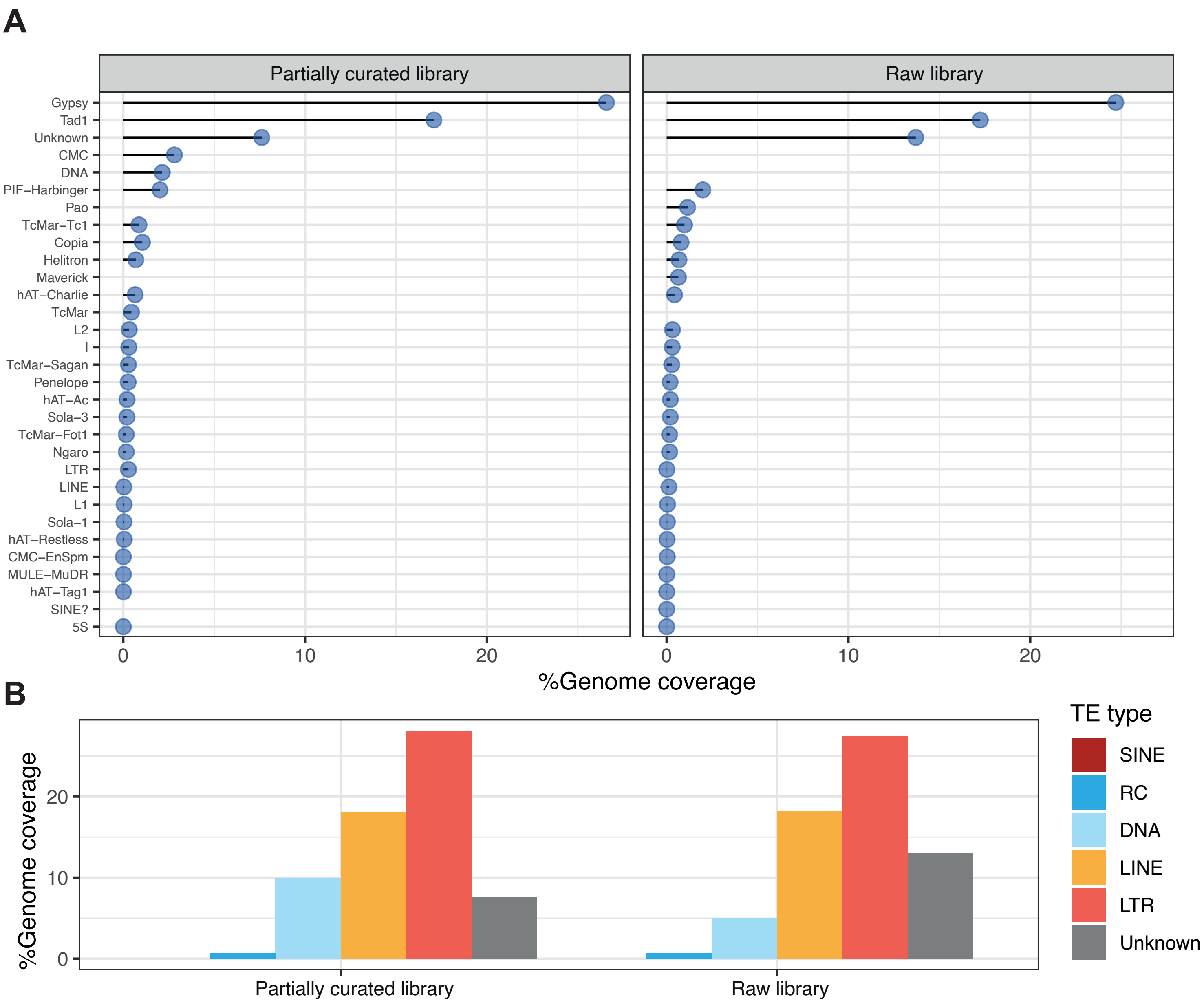


**Supplementary Figure S2**: Percentage of genome covered by (**A**) different transposon groups and (**B**) different TE class using the raw and the partially curated repeat libraries on the *T. panzihuhanense* genome.

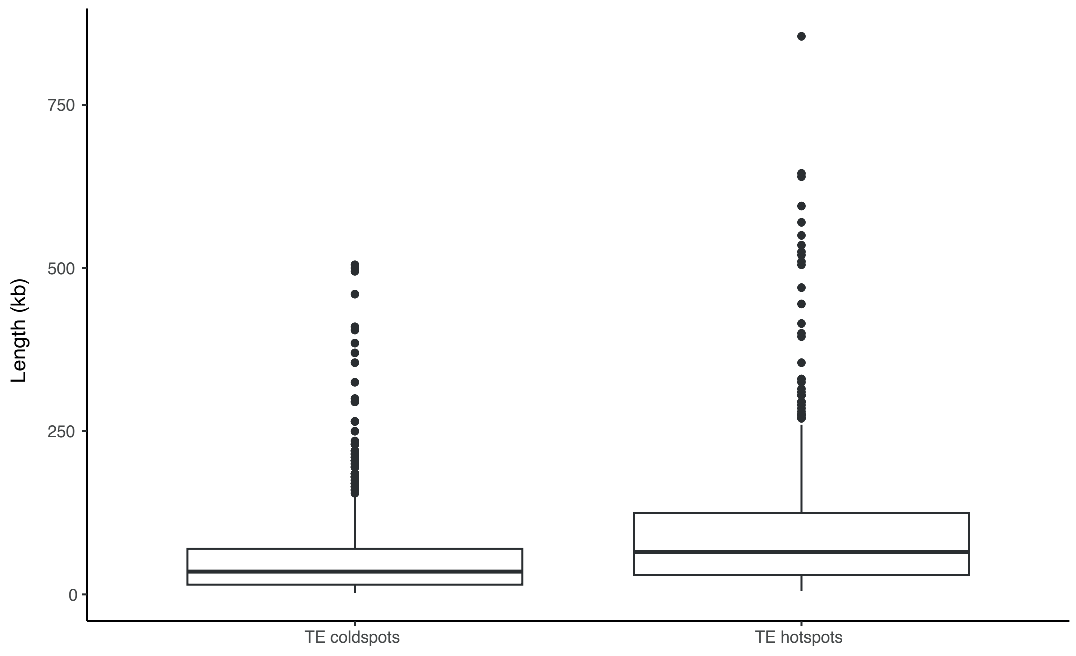


**Supplementary Figure S3**: ength in kilobases of significantly enriched (TE hotspots) and significantly depleted (TE coldspot) genomic regions in the *T. panzihuhanense* genome.

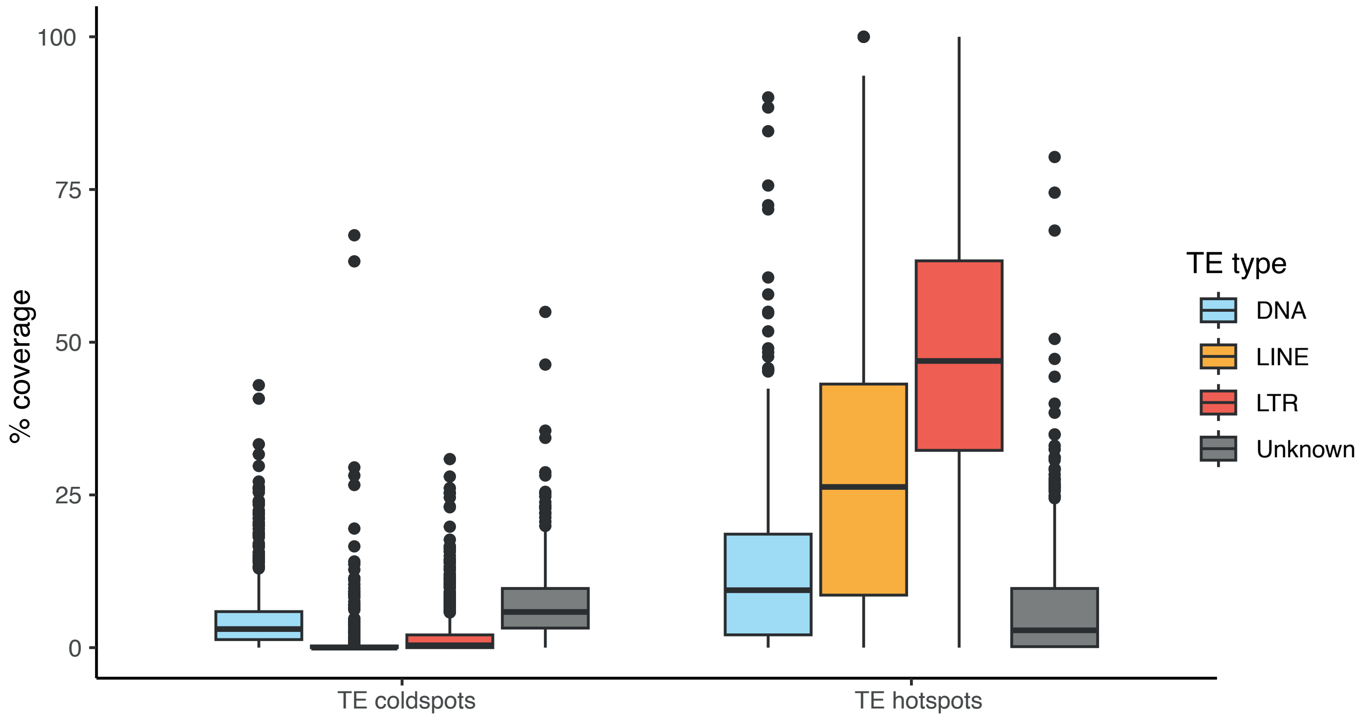


**Supplementary Figure S4**: Percentage of genomic coverage of the four most abundant transposon groups (DNA, LINEs, LTRs and Unknown) across TE coldspots and TE hotspots.

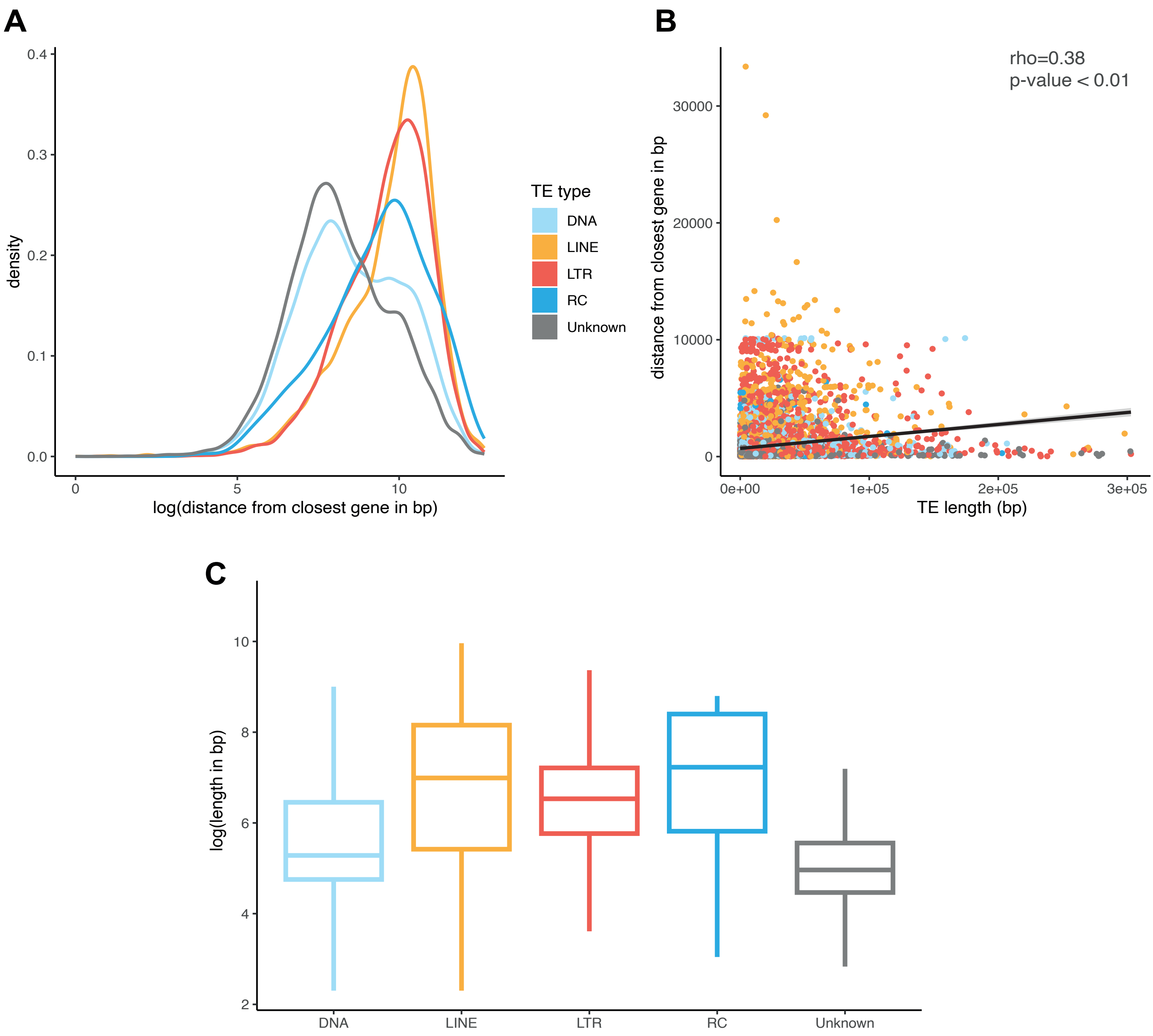


**Supplementary Figure S5**: (**A**) distance distribution from the closest gene for the 5 main transposon groups identified in the T. panzihuhanense genome. (**B**) Correlation between transposon length (y axis) and distance from the closest gene (x axis). (**C**) Length distribution of the 5 main transposon groups. In all plots Short Interspersed Nuclear Elements (SINEs) were excluded due to the low number of identified insertions.

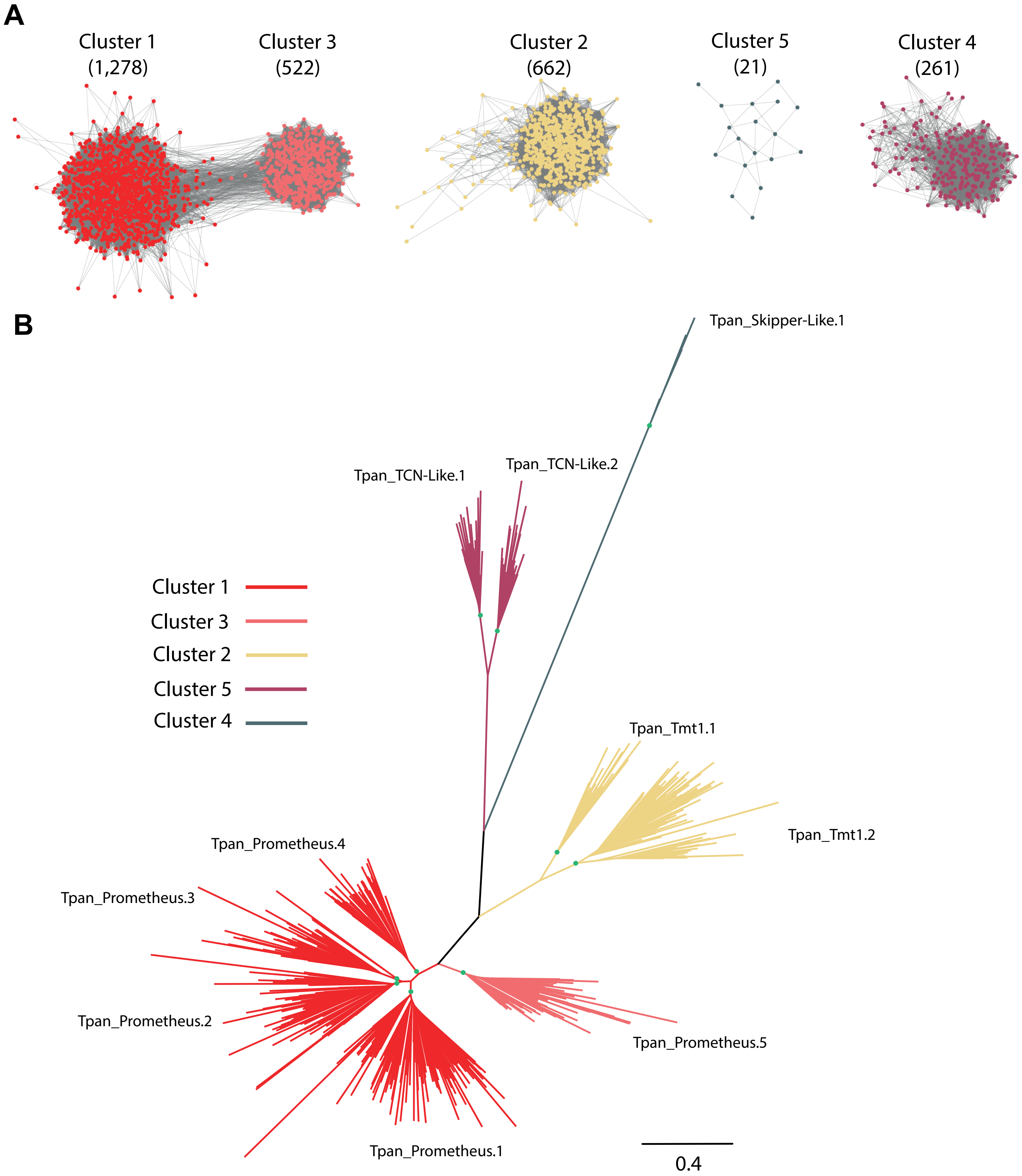


**Supplementary Figure S6:** Identification of the main long terminal repeats (LTR) Gypsy lineages in the T. panzihuhanense genome. (**A**) Communities identified through network analyses (using the greedy_modularity_communities algorithm in NetworkX) based on homologous relationships between reverse transcriptase (RT) nucleotide fragments mined from the T. panzihuhanense genome. Each cluster is labelled with its name, and the number of sequences in the respective cluster is shown in parentheses. (**B**) Phylogenetic relationships between representative RT protein segments. Colours correspond to those in panel A and represent clusters identified through network analyses. Families were identified based on cluster results, with some of them further split to highlight highly diverging, well-supported branches, represented by green circles (ultrafast bootstrap value ≥ 75).

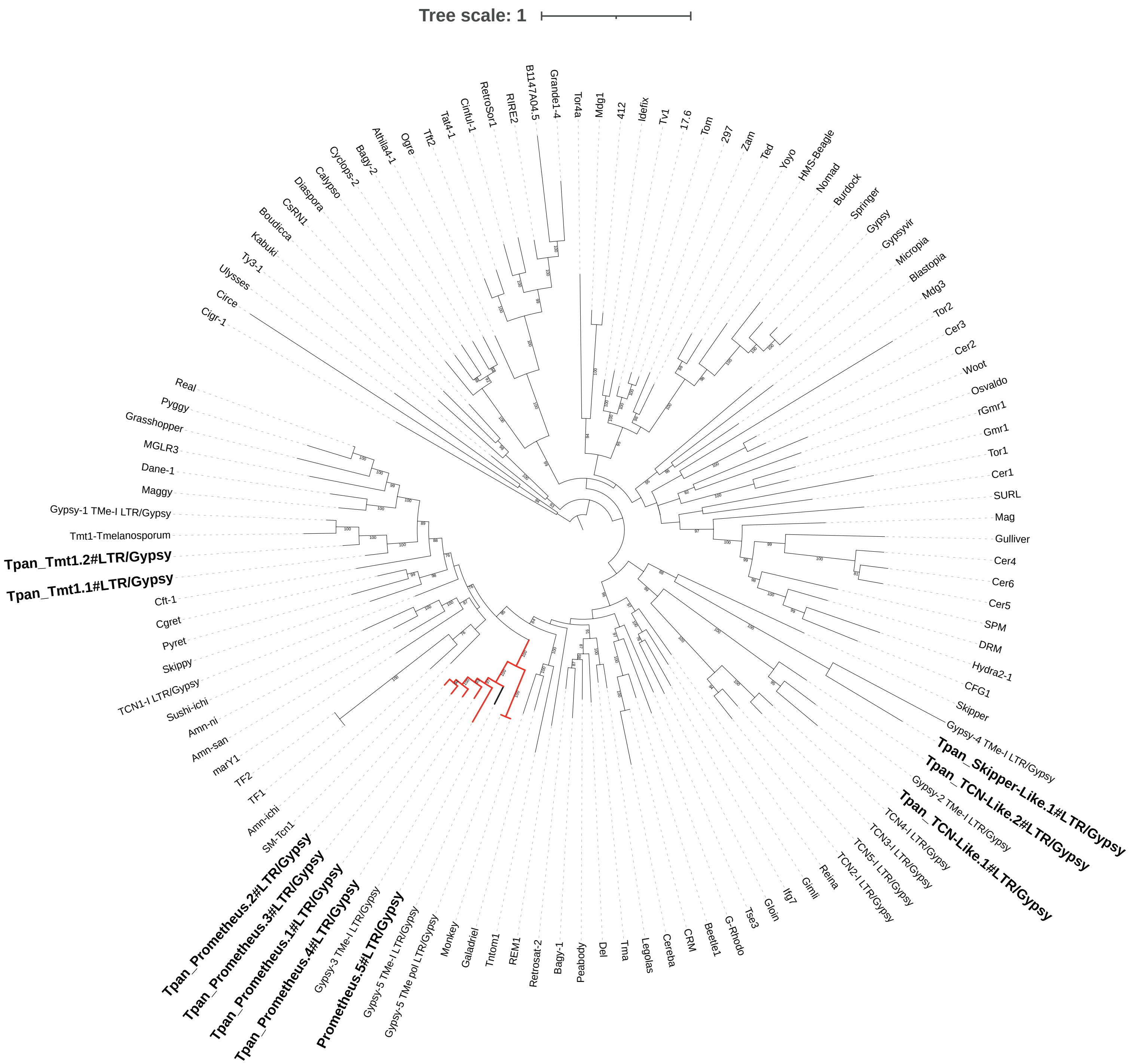


**Supplementary Figure S7**: Phylogenetic classification of *T. panzihuhanense* Gypsy families, highlighted in bold, based on reverse transcriptase domains and reference Gypsy sequences. Families highlighted in red represent the novel identified Prometheus clade. Numbers at each node represent ultrafast bootstrap values.

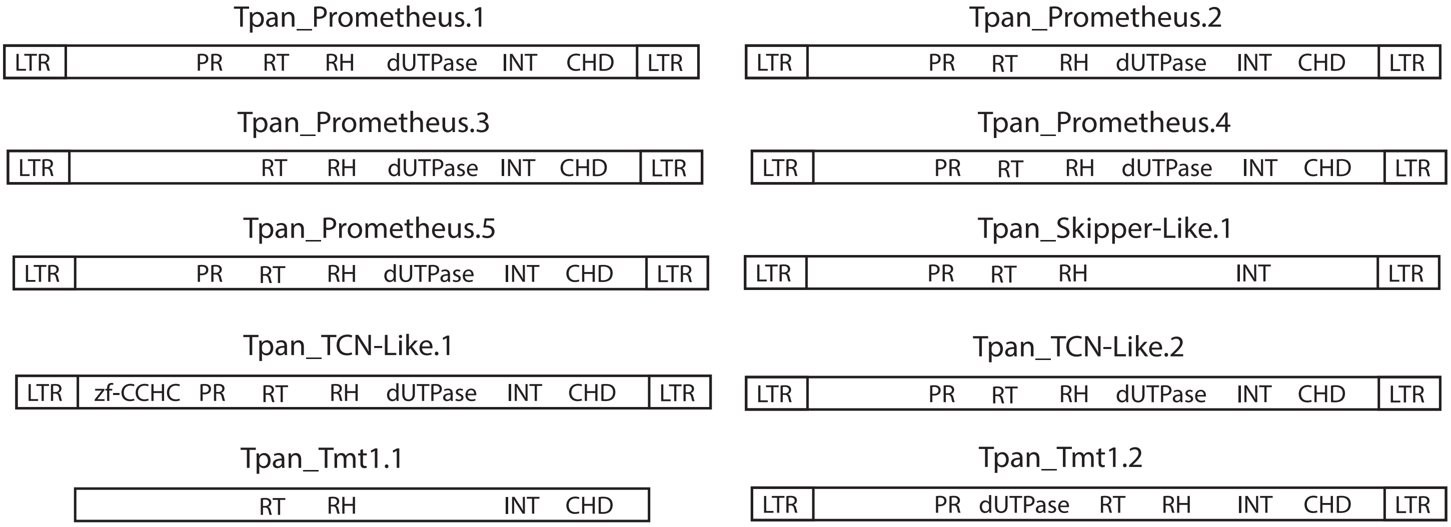


**Supplementary Figure S8**: Domain structure of reconstructed Gypsy families. For Tpan_Tmt1.1 we could not reconstruct a full length copy which included long terminal repeats segments. zf-CCHC = zinc knuckle binding motif; PR = proteinase; RT = Reverse transcriptase; RH = Ribonuclease H; dUTPase = Trimeric dUTP diphosphatases; INT = Integrase; CHD = Chromodomain; LTR = Long terminal repeats.

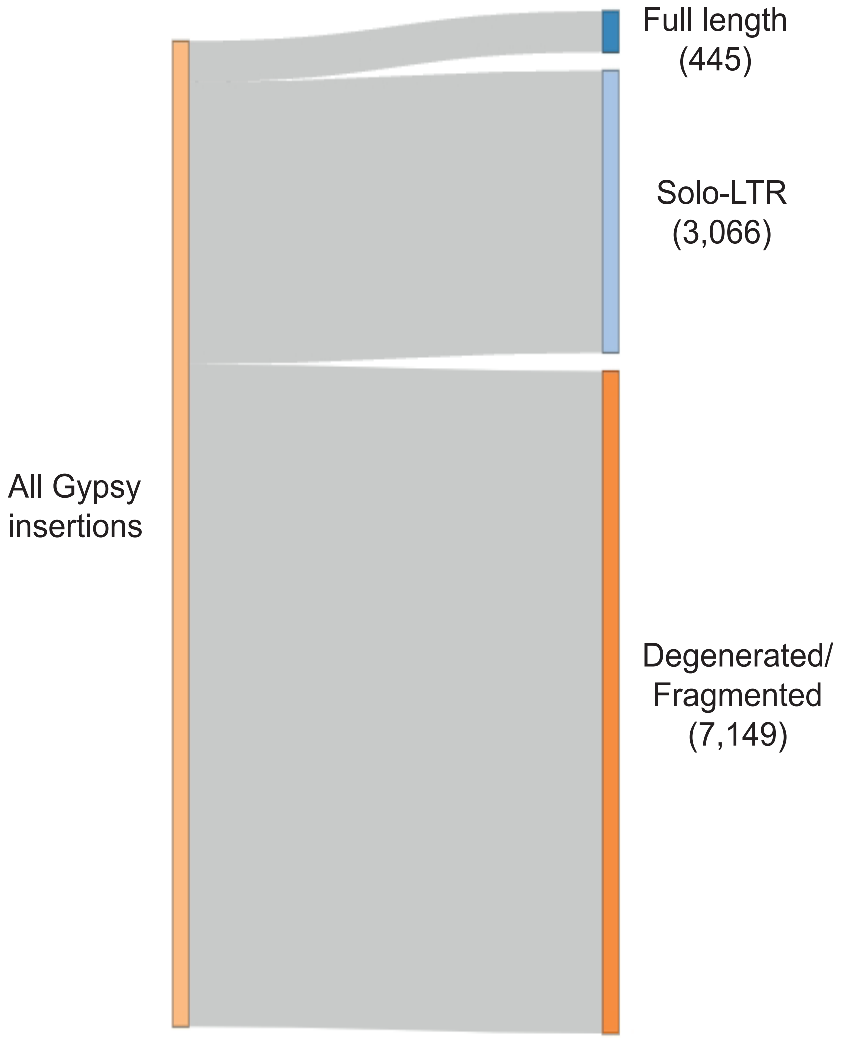


**Supplementary Figure S9**: Number of full length, solo-LTR and degenerated/fragmented Gypsy insertions identified in the T. panzhihuanense genome. Full length copies were defined as insertion covering the 90% of their parental consensus sequence. solo-LTR when both the insertion and on the two LTR segments of the parental consensus reciprocally aligned for at least 90%. All other instances were considered as degenerated/fragmented.

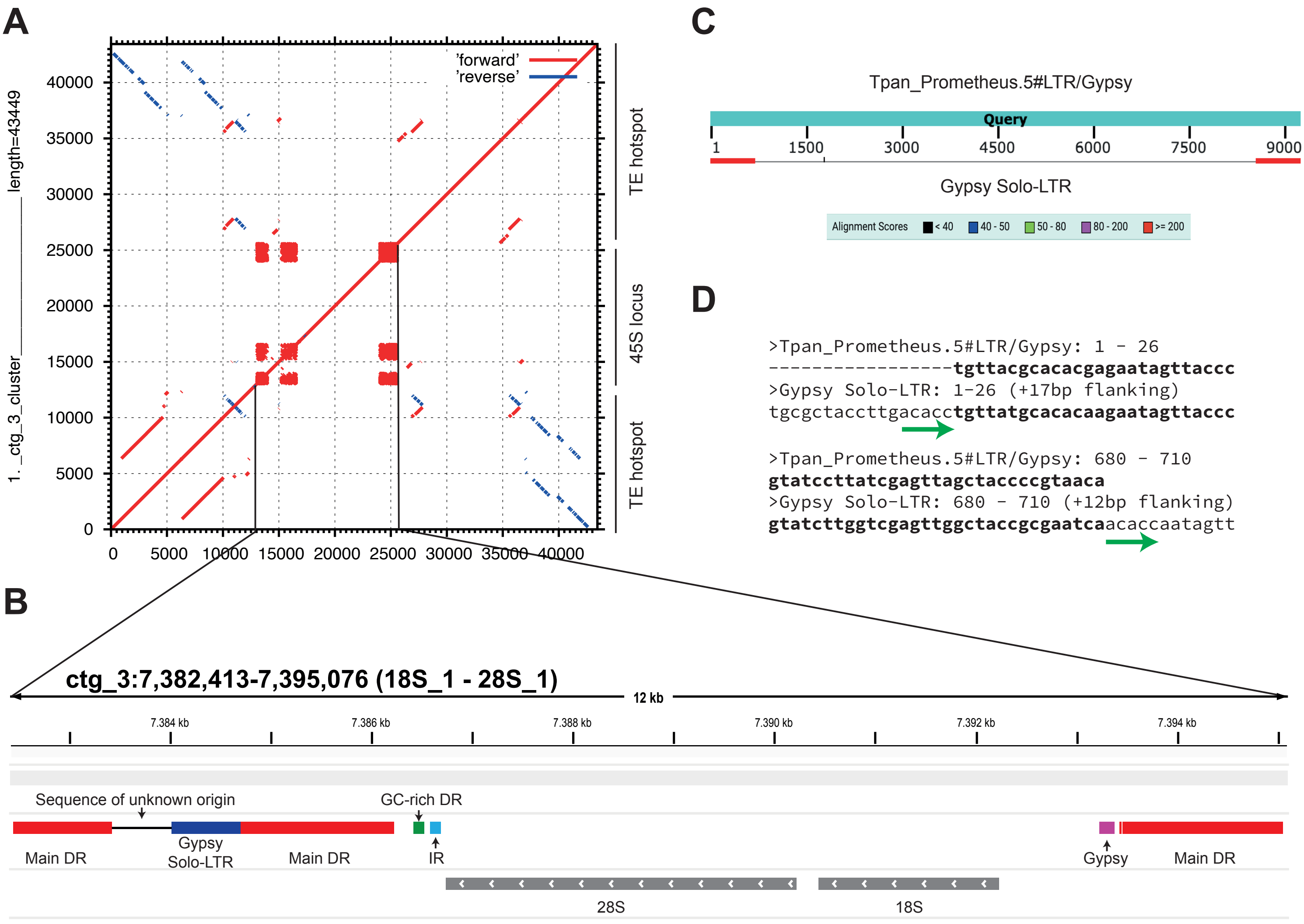


**Supplementary Figure S10**: Structure of the isolated 45S rDNA locus identified on ctg_3 and included in a complex genomic region corresponding to a TE-hotspot. (**A**) Self-alignment dot plot of the genomic region that includes the rDNA locus. Red lines represent alignments in the same orientation, while blue lines indicate alignments in the opposite orientation. (**B**) Detailed view of the structure and components of the rDNA locus. DR = direct repeats; IR = inverted repeats (**C**) BLASTn Alignment of the LTR fragment identified in the longer IGS region with its parental consensus sequence. (D) A more detailed view of the alignment, including the flanking regions of the solo-LTR insertion. Green arrows highlight a 5 bp target site duplication.

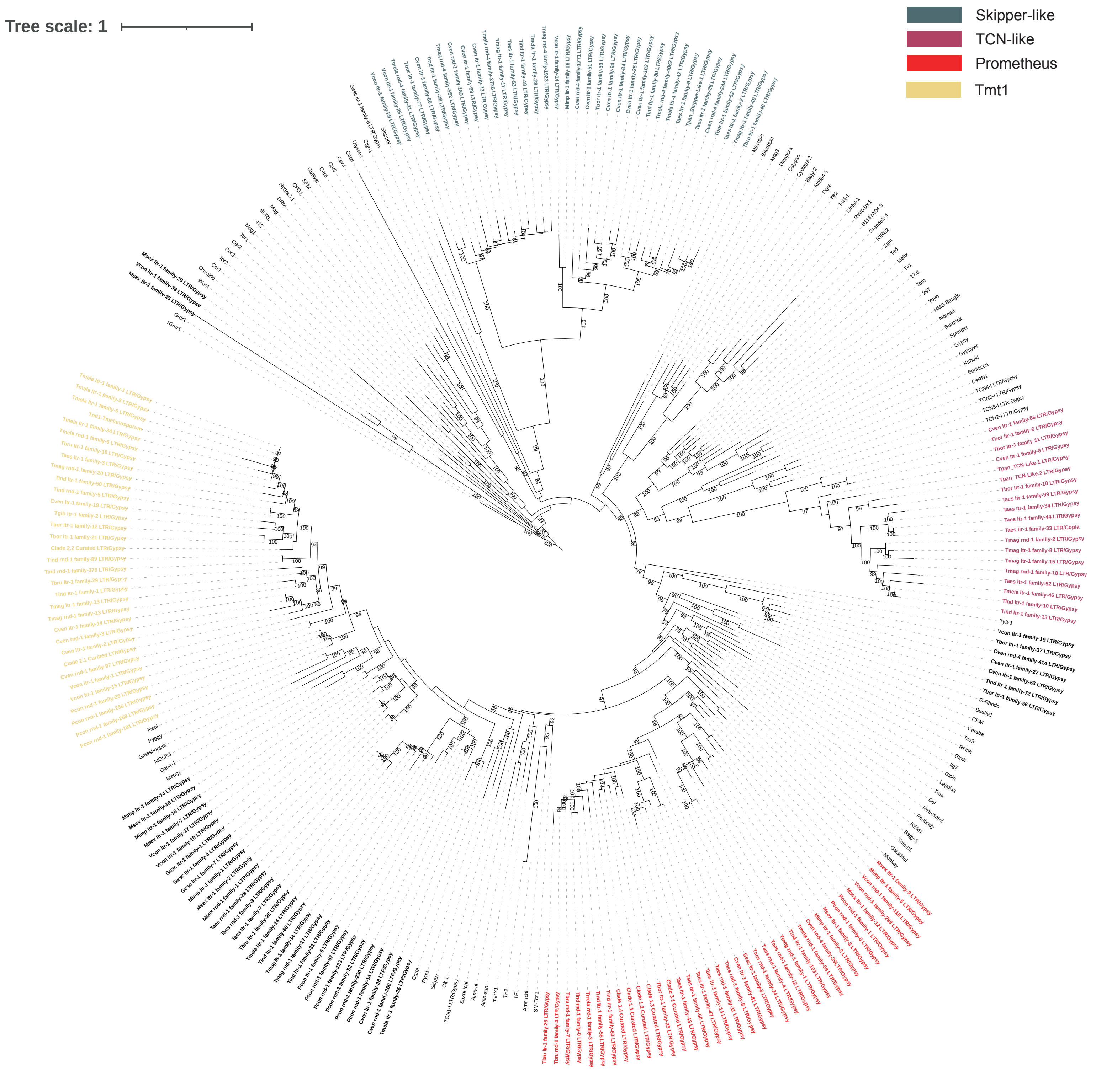


**Supplementary Figure S11**: Classification of Pezizales LTR Gypsy elements (in bold) based on phylogenetic relationships with known elements estimated on their reverse transcriptase protein domains. Different colours highlight the four clades identified also in T. panzhihuanense. Numbers at each node represent ultrafast bootstrap values. Each Pezizales element has a prefix reporting the source genome: Pcon = *Pyronema confluens*; Gesc = *Gyromitra esculenta*; Mimp = *Morchella importuna*; Msex = *Morchella sextelata*; Vcon = *Verpa coninca*; Cven = *Choiromyces venosus*; Tpan = *Tuber panzhihuanense*; Tbor = *Tuber borchii*; Tmag = *Tuber magnatum*; Taes = *Tuber aestivum*; Tbru = *Tuber brumale*; Tmela = *Tuber melanosporum*; Tind = *Tuber indicum.*

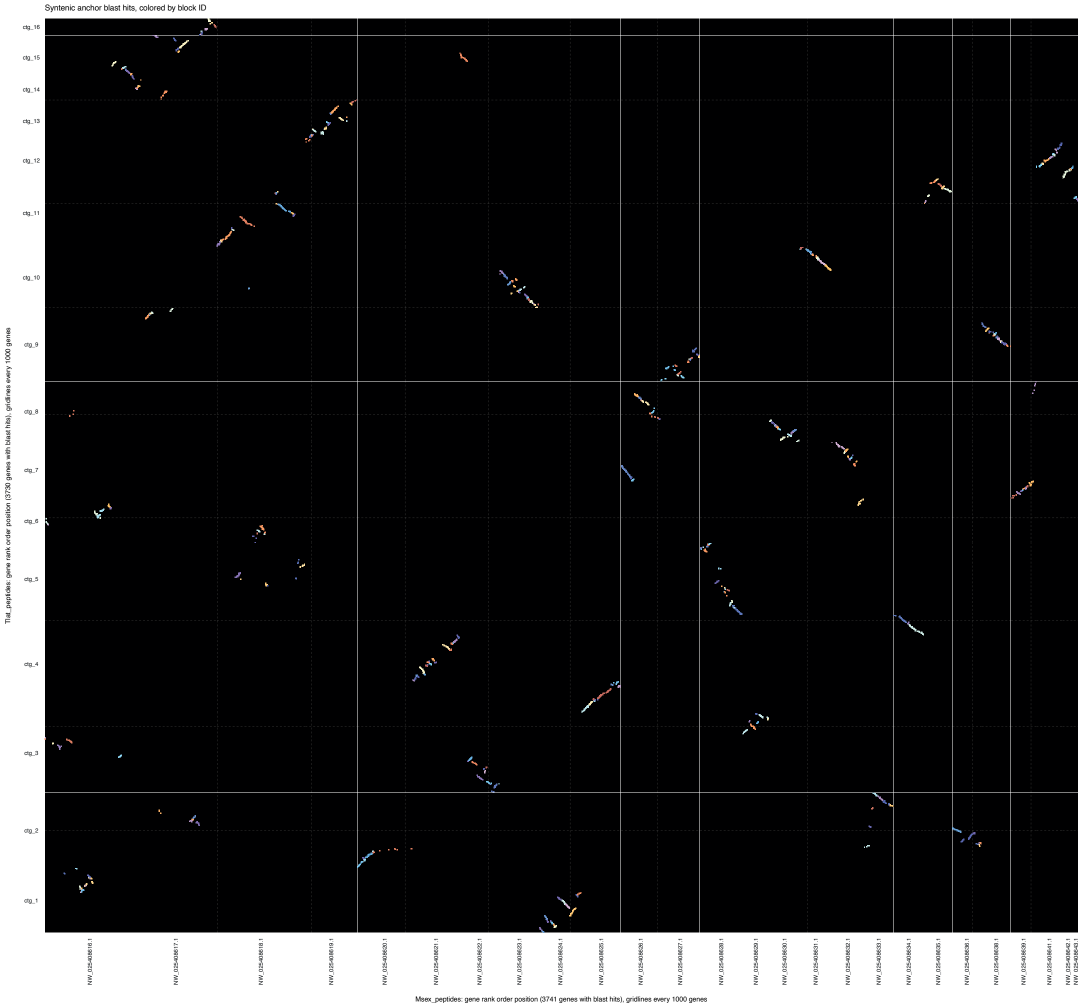


**Supplementary Figure S12**: Collinearity between *T. panzhihuanense* (y axis) and *M. sextelata* (x axis) genomes as inferred by GENESPACE.

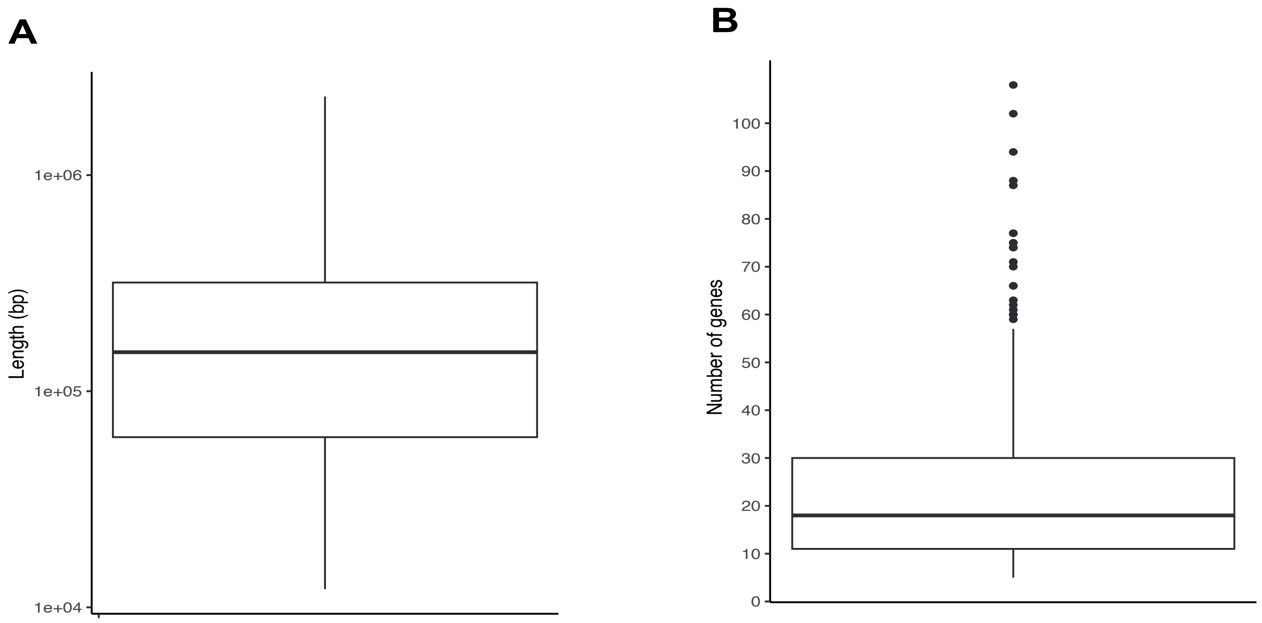


**Supplementary Figure S13**: (**A**) Length of syntenic blocks with *M. sextelata* in the *T. panzhihuanense* genome and (**B**) number of genes contained within these regions.

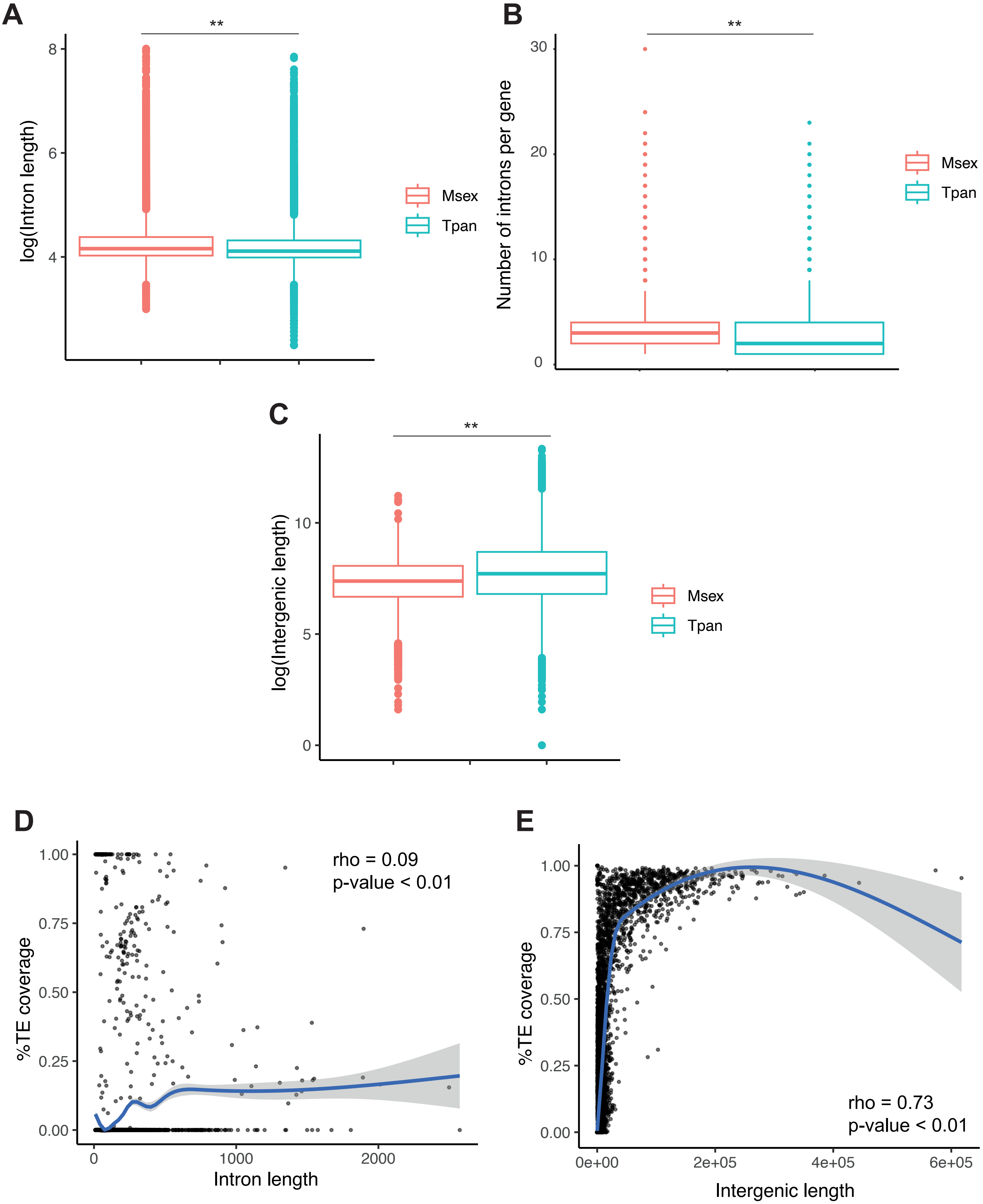


**Supplementary Figure S14**: Comparison between (**A**) intron length, (**B**) mean number of introns per gene and (**C**) intergenic genomic regions length between *T. panzhihuanense* (Tpan) and *M. sextelata* (Msex). (**D**) and (**E**) correlations between percentage of repeat coverage and intron and intergenic genomic regions length in T. panzhihuanense, respectively. rho = Spearman's rank correlation coefficient.

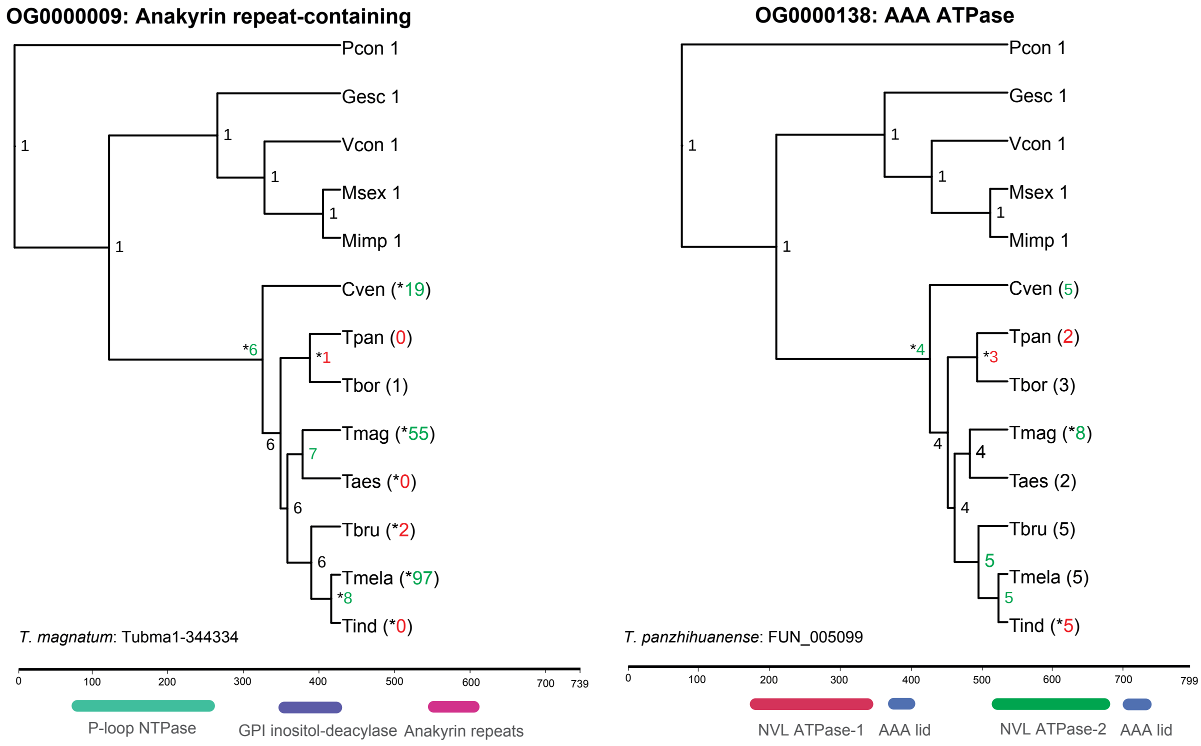


**Supplementary Figure S15**: CAFE results for the ECM-induced gene families OG0000009 and OG0000138 encoding for Ankaryn repeat containing proteins and AAA ATPAse, respectively. These families together with those represented in Fig. 6 were found significantly expanded in the stem branch of Tuberaceae. For each node, the inferred and observed gene family counts are reported for internal and terminal branches, respectively. Green colours highlight expansion events, red colours indicate contractions, and significant changes are marked with an asterisk. The domain annotation was obtained with InterProScan on a representative protein from T. magnatum. Pcon = Pyronema confluens; Gesc = *Gyromitra esculenta*; Mimp = *Morchella importuna*; Msex = *Morchella sextelata*; Vcon = *Verpa coninca*; Cven = *Choiromyces venosus*; Tpan = *Tuber panzhihuanense*; Tbor = *Tuber borchii*; Tmag = *Tuber magnatum*; Taes = *Tuber aestivum*; Tbru = *Tuber brumale*; Tmela = *Tuber melanosporum*; Tind = *Tuber indicum*.
