## Supplementary Material and Methods for "The high quality Chinese white truffle genome and novel fossil-calibrated estimate of Pezizomycetes divergence reveal the tempo and mode of true truffles genome evolution"

### Supplementary Materials and Methods

#### 1. Sampling and DNA sequencing

The sample used for genome sequences was the gleba tissue of a fruiting body (FB) of *T. panzhihuanense*. Morphological identification of this FB individual was made based on ascus and ascospore characteristics under a Leica CTR6-Leica DFC495-Leica DM6B system, compared with the description when the species of *T. panzhihuanense* was first discovered and classified from *T. latisporum* (Deng et al. 2013). Molecular characterization was confirmed after amplifying and sequencing the ITS region (Forward primer: ITS5 5'-GGAAGTAAAAGTCGTAACAAGG-3; reverse primer: ITS4 5'-TCCTCCGCTTATTGATATGC-3'; White et al. 1990). The rest of the FB was stored at −80°C while sending for sequencing. High-molecular-weight DNA extraction on the gleba was carried out through a modified CTAB protocol (patent number: ZL 2015 1 0617313.6), sheared to an average fragment size of ~20 kb using BluePippin Size-election system and HiFi SMRTbell library was constructed with the SMRTbell Express Template Prep Kit 1.0 (Pacific Biosciences). The same extracted DNA was used for the standard library construction process of Illumina TruSeq DNA PCR-free prep kit to produce a 400 bp insert size library.

#### 2. Sample preparation and RNA sequencing

*P. massoniana* seeds (Richu Seed Industry Co., Ltd., Suqian, China) that have been through cold stratification (at 4 ℃ for half a month) were sterilized in 30% $H_2O_2$ for 5 min before sowing in the seedling substrate [1:1 (v:v) perlite and vermiculite].

*T. panzhihuanense* fruiting bodies (FBs) used for making inoculum were purchased from Sichuan Golden Truffle Trading Co., Ltd. (Panzhihua, China) on Dec. 4th, 2020, with all individuals complying with Chinese GB/T38697-2020 standards. Molecular identification of the individual FB used for making inocula was carried out using species-specific primers TwitsF(5'-3'TAATGGTTTGAGGTGTTTGG)/TwitsR(5'-3'GTGAGATTTGCACTGTAAG).

Seedlings with basically the same growth status were selected 3 months after sowing. Each seedling was inoculated with 5 ml of a spore suspension containing $3\times10^{7}$ mature spores/ml.

Ascospore inoculation method was conducted, while the ascospore inocula was made of gleba tissues of *T. panzhihuanense* scattered in 5% sterilized gelatin. The concentration of inocula ($3\times10^7$ mature spores/ml) was set by a cell counting board (Qiujing Biochemical Reagent & Instrument, Shanghai), and inoculation was achieved by contact between inocula and roots of each seedling. The secondary inoculation was performed 3 months after the first one, with supplementary ascospores: the same dose of inocula prepared in the same way injected carefully by syringe into the mycorrhizal rhizosphere.

The whole process of ectomycorrhizal synthesis was carried out on a substrate obtained by mixing peat, quartz sand, perlite, vermiculite, and soil from natural truffle producing areas in the Jinsha River Basin (soil source location: 26.3847N 101.6331E, Chuxiong, China) at a ratio of 2:1:1:1:5 (v: v: v: v: v). Host seedlings grew in a light room that had been sterilized by potassium permanganate fumigation. 28W T8 #4 full-spectrum plant growth lights (Zhen'an Plant Growth Light Store, Xiamen) were used for lighting. The lighting intensity was set to 7500 lx according to the average light intensity on the sunny side of the mountainous area of Chuxiong Prefecture, Yunnan province, China (http://www.cxs.gov.cn/). The lighting time was set to 14 h per day. The indoor temperature was controlled at 25 °C, with humidity set to be 60% and the ventilation system running continuously. The water used for irrigation was sterilized nutrient solution: $Ca(NO_3)_2{\cdot}4H_2O$ 472 mg/L, $KNO_3$ 202 mg/L, $(NH_4)_2SO_4$ 80 mg/L, $KH_2PO_4$ 100 mg/L, $K_2SO_4$ 174 mg/L, $MgSO_4{\cdot}7H_2O$ 246 mg/L, $H_3BO_3$ 2.86 mg/L, $MnSO_4{\cdot}4H_2O$ 2.13 mg/L, $ZnSO_4{\cdot}7H_2O$ 0.22 mg/L, $CuSO_4{\cdot}5H_2O$ 0.08 mg/L, $(NH_4)_6Mo_7O_{24}{\cdot}4H_2O$ 0.02 mg/L.

6 months after the secondary inoculation, the ectomycorrhizae (ECMs) and the control samples (1-year-old uninoculated *P. massoniana* vegetative root tips) were collected.

Before RNA extraction, the identity of the selected mycorrhizas was molecularly identified to determine whether it was of the target species. DNA of 72 ectomycorrhizal root tips of each replicate seedling was extracted using the single nodule DNA extraction method for rhizobia (Terefework et al., 2001). Due to the low amount of DNA from ECMs, a two-round NEST-PCR was performed with primer pairs ITS1/4 and then TwitsF/R. Hosts whose ECMs produced 320bp bright bands then had another 100 ECMs (root tips for the control) taken as RNA sequencing materials. 3 biological replicates (as PMTW01 PMTW02 PMTW03, and PMCON01 PMCON02 PMCON03 for the control) were considered.

RNA integrity and the total amount have been assessed with the use of RNA Nano 6000 Assay Kit of Bioanalyzer 2100 system (Agilent technologies, CA, USA). mRNA has been isolated from total RNA with oligo(dT)-attached magnetic beads. Divalent cations were used under elevated temperature in Synthesis Reaction Buffer 5X for fragmentation. The first strand of cDNA has been synthesized through random hexamer primer and M-MuLV Reverse Transcriptase, followed by a second strand cDNA synthesis. Exonuclease/polymerase activities were applied to convert residual overhangs into blunt ends. As hybridization preparation, adaptors with hairpin loop structures were ligated following adenylation of the 3' ends of DNA fragments. cDNA fragments, preferably between 370 and 420 bp in length, were selected through library fragment purification using the AMPure XP system (Beckman Coulter, Beverly, USA). Subsequent PCR amplification was performed, and the PCR products were purified with AMPure XP beads, resulting in the final library. The library was quantified using the Qubit 2.0 Fluorometer, diluted to 1.5 ng/µl, and insert size was assessed with the Agilent 2100 Bioanalyzer. To accurately quantify the effective concentration of the library, qRT-PCR was employed, ensuring it exceeded 2 nM in order to maintain high quality.

### 3. Selection of fossils and calibration nodes for divergence time estimation

Previous estimates of truffle divergence based on genome data (Murat et al., 2018) have set the age of the most recent common ancestor (MRCA) of Pezizomycetes at circa 470 million years (MY) and the divergence of the Tuberaceae at circa 140 MYA. These dates have been estimated using as a calibration the posterior estimates for Tuberaceae MRCA (134 and 179 MYA) as defined by a previous work (Bonito et al. 2013). Bonito et al. in 2013, in turn, inferred divergence using the LSU substitution rate ($6.5\times10^{-4}$ substitutions per site per million years) defined on the divergences of the Erysiphales, a fungal order that is very different to the Pezizales both in terms of genetic distances and biology (Takamatsu and Matsuda 2004). Therefore, previous attempts to infer Pezizales divergence did not take into account fossils, which are more reliable than specific rates. For these reasons, we employed a multi-fossil calibration approach to estimate Pezizales divergences. We extensively inspected the mico-paleontological literature and defined various calibration points within the outgroups of the Pezizales. We calibrated the following nodes using the (range of) age of fossils as a minimum.

| Node | Fossil | Age (MYA) |
|---|---|---|
| Pleosporales (crown) | *Palaeocurvularia variabilis* | 95-93 |
| Aspergillus (crown) | *Aspergillus* sp. | 55-35 |
| Parmelia - Xanthoparmelia split | Parmelioid fossils | 45-15 |
| Ophiocordyceps (crown) | *Paleoophiocordyceps coccophagus* | 105-99 |
| Glomerellaceae (crown) | *Colletotrichum* sp. | 67.8–61.6 |
| Botrytis - Cyclaneusma split | *Ovularites barbouri* | 145.5-65.5 |
| Arthrobotrys (crown) | *Palaeoanellus dimorphus* | 113-100.5 |
| Pezizales (crown) | *Fusiformisporites acutus* | 23.3-11.6 |
| Pezizomycotina (stem) | LL03/02 NMW 2018.17G.10 | 425.6-423.0 |

The reasons behind the choice of fossils are discussed in the following paragraphs.

Ascomycota and Pezizomycotina. *Paleopyrenomycites devonicus* is an extinct species belonging to the subkingdom Dikarya, the group that includes the phyla Ascomycota and Basidiomycota. It was discovered in the Rhynie Chert, 407 MYA (Taylor et al. 1999; Taylor et al. 2005). It is one of the oldest known representatives of the Dikarya, and the only one with both sexual and asexual morphs preserved in the same specimen. The question of whether it occupies a basal position within the Ascomycota or is nested within the subphylum Pezizomycotina remains a subject of debate. Its precise placement varies depending on the specific interpretation, resulting in significantly divergent age estimates for the same divergence events (Taylor and Berbee 2006; Lücking et al. 2009; Prieto and Wedin 2013; Beimforde et al. 2014; Lutzoni et al. 2018). Strullu-Derrien et al. (2023) suggested using *P. devonicus* as a minimum node age calibration point for the entire Ascomycota phylum, potentially representing the stem node of Ascomycota or even the Dikarya crown group. The systematic position of *Prototaxites* fossils (Upper Devonian, Famennian stage) has been longly debated; it has been assigned to bryophyte-grade streptophytes, chlorophytan green algae, rhodophytan red algae, phaeophycean brown algae, and fungi belonging to three phyla (Basidiomycota, Mucoromycota, Ascomycota). Finally, after an extensive analysis (Hueber 2001), it has been largely accepted as belonging to the kingdom Fungi (Strullu-Derrien et al. 2023). A recent work assigned the fossil species *P. taiti* to a basal Ascomycete belonging to Neolectomycetes, Taphrinomycotina or Pezizomycotina (Honegger et al. 2018). Strullu-Derrien et al. (2023) suggest using the hymenial layer bearing polysporus asci attributed to *P. taiti* to be used as a calibration for the stem node in Ascomycota. *P. taiti* is found in the Pragian Rhynie chert, Aberdeenshire, from lower Devonian, dating 407 MYA. However, a charcoalified, unnamed Upper Silurian (upper Ludlow, 423.0-425.6 MYA) apothecial fragment LL03/02, as well as the NMW 2018.17G.10 fragment, discovered in Ludford Lane, Shropshire, exhibit characteristics indicative of their placement within Pezizomycotina (Edwards et al. 2018; Honegger et al. 2018). Thus, we decided to use the unnamed Pezizomycotina (instead of the updated *Paleopyrenomycites devonicus* and *Prototaxites taiti)* as a minimum for the Pezizomycotina stem (423 MYA). We set a root prior for the crown Ascomycota (split of Taphrinomycetes) using a normal distribution centred at 534.0 MYA, allowing a permissive sampling between 497.6 and 570.8 MYA; this is in accordance with the comparison of 17 posterior distributions as defined by TimeTree (Kumar et al. 2022).

Pezizomycetes. *Pezizasporites taiwanensis* is characterised by elliptical ascospores from the Miocene of Taiwan that are up to 20 µm long (Huang 1981). It is thought to belong to the Pezizales order (Saxena 2021). *Fusiformisporites acutus*, from the Lower-Middle Miocene (Kumar 1990), seems to belong to the Sarcoscyphaceae family, order Pezizales (Saxena 2021). We decided to use the age of the Lower Miocene to calibrate the Pezizales crown group (split between *Tuber* and *Ascobulus immersus*) minimum (23.3).

Orbiliomycetes. Among the Orbiliomycetes, *Palaeoanellus dimorphus* is a predatory fungus from the late Albian (Schmidt et al. 2007). It has been identified as an ancestor of the extant species *Arthrobotrys blastospora* (Zhang et al. 2023). It is a very useful calibration as it can be assigned at the genus level, but it has not yet been used in molecular clock analyses. We decided to use the upper bound of the Albian (100.5 MYA) as a minimum calibration of the *Arthrobotrys* crown (*A. flagrans* and *A. iridis* split).

Leotiomycetes. *Ovularites barbouri* is a species from the Cretaceous belonging to the Helotiales order (Whitford 1916). Although it stands as the sole representative of the Leotiomycetes class, it has yet to be employed in molecular clock studies. We used the upper boundary of the Cretaceous (66 MYA) as a minimum for the split of *Botrytis cinerea* and *Cyclaneusma minus*.

Sordariomycetes. The class Sordariomycetes was calibrated by Lutzoni et al. in 2018 with a *Colletotrichum* fossil in Burmese amber found in Isisaurus dung from the upper Cretaceous. Simil fossils are present in the intertrappean Beds from the late Cretaceous, dating 67.8–61.6 MYA (Kar et al. 2004; Kar and Verma 2004); since *Colletotrichum* is the anamorph of *Glomerella* (monotypic family Glomerellaceae, Glomerellales, Sordariomycetes). We calibrate the crown group of *Glomerellaceae (*split between *Colletotrichum gloeosporioides* and *Colletotrichum destructivum)* using, as a minimum, the upper late Cretaceous age (61.6). Sung et al. in 2008 described a new fossil, *Paleoophiocordyceps coccophagus,* an asexual state of the genus *Ophiocordyceps*. The fossil was found in Burmese amber from the upper Albian (Early Cretaceous, 99–105 MYA) (Sung et al. 2008). We used it to calibrate the Ophiocordyceps crown (split of *Ophiocordyceps australis* and *Ophiocordyceps sinensis*) with a minimum of 99 MYA.

Lecanoromycetes. The parmelioid fossil *Alectoria succinica* (Mägdefrau 1957*)*, previously employed as a calibration point (Amo de Paz et al. 2011; Prieto and Wedin 2013), has been found to lack morphological features typical of alectorioid lichens. Instead, it appears to be a degraded plant part, likely a root (Kaasalainen et al. 2015). Therefore, we decided not to use it as a calibration point. Two other parmelioid species have been described from Dominican amber (Poinar Jr et al. 2000) dating 15–45 MYA; they have been used to calibrate the crown of the *Parmelia* genus (Poinar Jr et al. 2000; Amo de Paz et al. 2011), the split of the Parmelia–Alectoria clade (Prieto and Wedin 2013) or the Parmeliaceae family (Del-Prado et al. 2013) on the basis of authors' interpretation. Given that the genus *Parmelia* has a singular genome representation, this calibration point would only serve as a conservative estimate for the divergence time between *Parmelia* and its closest relative with a sequenced genome, *Xanthoparmelia taractica*; therefore we used the upper age of the amber (15 MYA) as a minimum for the split of *Parmelia* sp. and *Xanthoparmelia taractica*.

Dothideomycetes. An *Aspergillus* (Aspergillaceae, Eurotiales) fossil (Dörfelt and Schmidt 2005) was found in Baltic amber (55–35 MYA) associated with collembola species. Following the approach of Beimforde et al. in 2014, who employed it as a minimum prior for the oldest *Aspergillus* species, we used it to calibrate the *Aspergillus* crown using the amber minimum age (35 MYA) for the split of *Aspergillus wentii* and *Aspergillus niger*. *Palaeocurvularia variabilis* is one of the oldest representatives of the Pleosporales order. It was found in Ethiopian amber, dating back to 95-93 MYA (Schmidt et al. 2010). It was used to calibrate the split between Dothideomycetes and Arthoniomycetes (Gueidan et al. 2011), while Beimforde et al. in 2014 decided to remove it from the analysis due to its lack of assignment to any modern family. We decided to use it as a minimum calibration (93 MYA) of the Pleosporales crown group by calibrating the split of Cucurbitariaceae and Delitschiaceae.

Other fungal fossil calibrations identified but not included in this analysis because of a lack of genomes and/or to avoid too many taxa and consequent computational issues are listed below.

Lecanoromycetes. Fossils attributed to the Lecanoromycetes class have been categorised into two orders: Lecanorales and Caliciales. The order Lecanorales is primarily represented in the fossil record by the family Parmeliaceae. Among these fossils, *Anzia electra*, discovered in Baltic amber, dates back 35-40 million years. This specimen readily fits into the genus *Anzia*, specifically within the *Anzia* sect *Anzia* (Rikkinen and Poinar Jr 2002). Lutzoni et al. utilised this fossil as a minimum age constraint for the Parmelioid clade in 2018, while Beimborde et al. in 2014 utilised it as a calibration for the *Anzia*-*Canoparmelia* split. However, due to the absence of genomic data for this genus, we have not used it in our analysis. *Phyllopsora dominicanus*, a fossil discovered in Dominican amber by Rikkinen and Poinar Jr in 2008, is classified within the Ramaliaceae family. However, it was excluded from the analysis conducted by Beimforde et al. in 2014 due to redundancy resulting from the presence of other fossils belonging to the Lecanorales clade. Within the order Caliciales, a solitary fossil named *Calicium succini*, belonging to the family Caliciaceae, has been identified. This specimen was unearthed in Baltic amber, dating back 24-47 million years ago. The preserved characteristics closely resemble those observed in present-day species of *Calicium* (Rikkinen et al. 2018). However, the absence of a reference genome for the genus *Calicium*, as well as for the entire Caliciaceae family, restricts its utility to serving solely as a calibration point for the crown group of the Caliciales order and we did not use it.

Dothideomycetes. *Pteropus brachyphylli* is a fossil found in silicified conifer leaves from the Valkenburg Member (Maastricht Formation), Belgium, dating back 66.5 MYA. It was thought to belong to the Venturiaceae family (van der Ham and Dortangs 2005). However, its uncertain taxonomic placement raises questions about the corresponding modern genus *Phaeocryptopus*, which is likely polyphyletic and may belong to either the Dothideales or Capnodiales and, therefore, not used for calibration for precaution (Beimforde et al. 2014): we followed this indication and excluded it from our analysis. A more definitive example within the Capnodiales is *Metacapnodium succinum*, exhibiting morphology nearly identical to extant *Metacapnodium* species. This specimen was discovered in European Amber dating back to 22-54 MYA (Rikkinen et al. 2003). Along with the older representative of Metacapnodiaceae (Schmidt et al. 2014) from the Albian stage, it was used to calibrate stem (Beimforde et al. 2014; Pérez-Ortega et al. 2016; Liu et al. 2017; Lutzoni et al. 2018) or crown (Hongsanan 2020) of the Metacapnodiaceae family. However, since there are no genomes representative of Metacapnodiaceae, we decided to exclude it from the analysis. The order Asterinales is represented in fossils with 9 species of Asterina. The oldest of them is *Asterina eocenica*, dating back 59–54 MYA (Dilcher 1965). It has been suggested that it be used to calibrate the crown node of Asterina as a minimum prior (Samarakoon et al. 2019). However, since there are no genomes representative of the order Asterinales, we could not use it. Among the Mycocaliciales, the fossils *Chaenothecopsis bitterfeldensis* (Rikkinen and Poinar 2000) dating 22-40 MYA and *Phaeocalicium* sp. (Rikkinen et al. 2018) of 23.8-25.3 MYA, were used to calibrate the order (Prieto and Wedin 2013; Beimforde et al. 2014; Pérez-Ortega et al. 2016) however, due to the absence of genomes for Mycocaliciales, we made the decision to exclude them from the analysis. *Asterothyrites* is a genus with nine described fossils, representative of the family Microthyriaceae, the sole family in the order Microthyriales. The oldest of which (*Asterothyrites dictyozamiticola*) is from the early Cretaceous (145.5–99.6). It has been suggested as a calibration point of the crown of Microthyriales (Samarakoon et al. 2019). Since there is only one genome in this family (*Microthyrium microscopicum*) we decided not to use it.

Coniocybomycetes. The calicioid fossil *Chaenotheca succina* was found in Paleogene Baltic amber dating back 38–34 MYA (Rikkinen et al. 2018) and used in molecular clock studies to calibrate the Coniocybomycetes class (Pérez-Ortega et al. 2016) or the Coniocybaceae family (Prieto and Wedin 2013) while Beimforde et al. 2014 decided to discard this fossil due to concerns about the potential cross-prior influence on age estimates, which could lead to bimodal posterior distributions in other constraints. Even if a recent study proposed utilising it as a minimum prior for the *Chaenotheca* genus (Samarakoon et al. 2019), we opted to adhere to Beimforde et al. recommendation in 2014.

Sordariomycetes. The fossil *Spataporthe taylori* dating 132.9 MYA at the Early Cretaceous Valanginian-Hauterivian boundary has morphological features belonging to the order Diaporthales, with characteristic ascus tips of the Gnomoniaceae family (Bronson et al. 2013). It was used to calibrate the stem node of the order Diaportales by (Guterres et al. 2018); however, it was not used in the study of (Chen et al. 2023) because a preliminary analysis showed that the obtained divergence times were older than the previous studies. We chose to follow this recommendation and disregard it.
